# Intra-epithelial non-canonical Activin A signalling safeguards prostate progenitor quiescence

**DOI:** 10.1101/2021.03.05.433921

**Authors:** Francesco Cambuli, Veronica Foletto, Alessandro Alaimo, Dario De Felice, Francesco Gandolfi, Maria Dilia Palumbieri, Michela Zaffagni, Sacha Genovesi, Marco Lorenzoni, Martina Celotti, Emiliana Bertossio, Giosuè Mazzero, Arianna Bertossi, Alessandra Bisio, Francesco Berardinelli, Antonio Antoccia, Marco Gaspari, Mattia Barbareschi, Michelangelo Fiorentino, Michael M. Shen, Massimo Loda, Alessandro Romanel, Andrea Lunardi

## Abstract

The healthy prostate is a relatively quiescent tissue. Yet, prostate epithelium overgrowth is a common condition during ageing, associated with urinary dysfunction and tumorigenesis. For over thirty years, TGF-β ligands have been known to induce cytostasis in a large variety of epithelia, but the intracellular pathway mediating this signal in the prostate, as well as its relevance for quiescence, have remained elusive.

Here, using mouse prostate organoids to model epithelial progenitors, we found that intra-epithelial non-canonical Activin A signalling inhibited cell proliferation in a Smad-independent manner. Mechanistically, Activin A triggered Tak1 and p38 MAPK activity, leading to p16 and p21 nuclear import. Spontaneous evasion from this quiescent state occurred upon prolonged culture, due to reduced Activin A secretion, a condition associated with DNA replication stress and aneuploidy. Organoids capable to escape quiescence *in vitro* were also able to implant with increased frequency into immunocompetent mice.

Our study demonstrates that non-canonical Activin A signalling safeguards epithelial quiescence in the healthy prostate, with potential implications for the understanding of cancer initiation, and the development of therapies targeting quiescent tumour progenitors.

## Introduction

The healthy prostate is a relatively quiescent tissue during adulthood^1,2^. In contrast, the overgrowth of the prostatic epithelium is one of the most common conditions experienced by ageing men, being linked with urinary dysfunction and tumorigenesis^3^. The molecular mechanisms causing exit from quiescence are poorly understood. Chronic inflammation – potentially induced by infection (*e.g.*, prostatitis)^4–6^, chemical damage (*e.g.*, urine reflux)^7^, physical trauma (*e.g.*, corpora amylacea)^8^, dietary carcinogens^9^, obesity^10^, hormonal imbalance (*e.g*., low systemic androgen levels)^11,12^, and ageing^13^ – has been implicated in DNA damage, oxidative stress, and atrophy, leading to a proliferative response^14,15^. Considering the high frequency of these events, it would be logical to hypothesize specialized mechanisms to safeguard epithelial quiescence, but they have been rarely investigated.

It has long been known that Transforming Growth Factor β (TGF-β) signalling inhibits the proliferation of a large variety of epithelial cell types^16,17^, including those of the prostate^18^. SMAD factors are the canonical intracellular mediators of this signalling, but additional non-canonical pathways can also be triggered by TGF-β receptors^19,20^. In gastrointestinal (GI) carcinomas (*e.g*., pancreas, colon), the canonical pathway is frequently mutated^21,22^. However, outside of the GI tract, TGF-β/SMAD components are rarely inactivated in tumours, leaving unexplained the nature of the intracellular signalling responsible for the cytostatic effect of TGF-β^23,24^.

Enhanced Tgf-β signalling has been linked with the presence of quiescent epithelial progenitors in the proximal/periurethral region of the mouse prostate^25,26^. Recent single-cell studies have confirmed the enrichment of a variety of epithelial progenitors – basal, luminal proximal (LumP), and periurethral (PrU) cells – in this anatomical district, though also present at low frequency in the distal compartment^27–32^. Such cells are known to be particularly quiescent during homeostasis^33,34^, but also to exhibit extensive regenerative potential in *ex-vivo* assays^31,35^.

Thus, the TGF-β induced cytostatic response in epithelial progenitors may be relevant for the control of quiescence, but the complexity of this pathway, the lack of interpretable genetic alterations in patients, and the heterogeneous cellular composition of the prostate, have so far hampered mechanistic investigations. Here, we reasoned that prostate organoid models^36,37^ – in combination with orthotopic transplantation approaches – may provide a biologically relevant, and experimentally amenable, system for addressing this question.

## Results

### Mouse prostate organoid cultures enable the continuous expansion of epithelial progenitors in a near-physiological manner

Initially, we set out to assess whether mouse prostate organoids are a representative and informative model for the study of the prostate epithelium, in light of recent discoveries on prostate cellular heterogeneity and dynamics^27,28,30-32,34,38^. Considering our interest in signalling, we focused on a culture method in defined media conditions. This protocol relies on a mix of growth factors and inhibitors, including Egf, Noggin, R-spondin 1, the Tgf-β receptors inhibitor A83-01, and dihydrotestosterone (ENRAD)^36^. We generated a biobank of mouse prostate organoids, starting from bulk populations of cells from distinct prostate lobes and mouse strains (Fig. 1a, Supplementary Fig S1). In line with previous studies^36,37^, we found that, upon tissue dissociation, only a small fraction of cells (approx. 1%) was capable to generate organoids in culture, and that organoid-forming efficiency increased over passages, suggesting enrichment for epithelial progenitors (Fig. 1b). To gain greater insights, we longitudinally tracked organoid formation – from single cells to fully formed organoids – and we observed a progressive expansion of cells expressing the progenitor epithelial surface antigen Sca-1 (encoded by *Ly6a*) (Fig. 1c)^31,35^. Thereafter, the level of Sca-1 appeared to be stable over a long culture period (*e.g*., 10 weeks). To extend our observations, we performed transcriptomic analyses on three organoid lines derived from distinct mouse prostate lobes (Fig. 1d,e). Consistently with enrichment for epithelial progenitors, organoids expressed high levels of genes specific for the proximal and periurethral compartments (*e.g*., *Psca, Tacstd2*, and *Ly6d*), as well as basal (*e.g*., *Krt5, Krt14, Trp63*) and luminal marker genes (*e.g., Krt8, Ar, Foxa1*). In contrast, distal luminal markers were barely detectable (*e.g., Nkx3.1, Pbsn, Sbp*). Histological (H&E) and immunofluorescent (IF) analyses confirmed that prostate organoids, for the most part, are made up of a bilayer of cuboidal cells, displaying progenitor marker proteins (*e.g*., Krt7, Ppp1r1b), and resembling the cyto-architecture of the periurethral/proximal compartment (Fig. 1f-h). As expected for periurethral/proximal cells – which are known to be castration resistant – mouse prostate organoids were reversibly dependent on androgen for lumen formation, but not for their survival (Supplementary Fig. S2).

**Figure 1.**
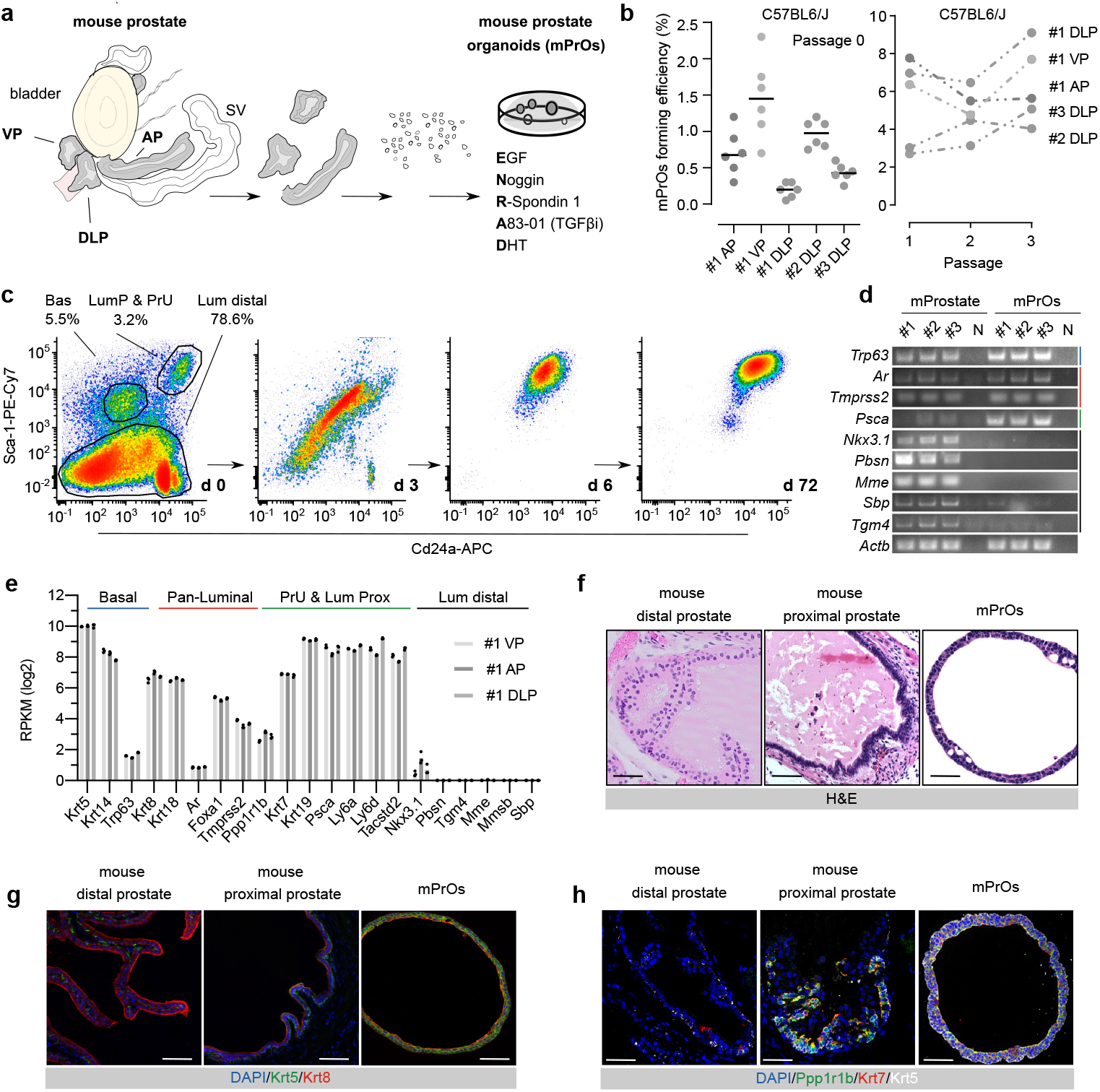
Mouse prostate organoids are highly enriched in epithelial progenitor cells. **a.** Schematic diagram describing organoid culture derivation (AP, Anterior Prostate; DLP, Dorso-Lateral Prostate; VP, Ventral Prostate; SV, Seminal Vesicle; DHT, dihydrotestosterone). **b.** Mouse prostate organoid forming efficiency. Efficiency at derivation (left; data points are shown with crossing line representing mean value). Efficiency at passage 1-3 (right; n≥ 3 per organoid line/passage; data are presented as mean). **c.** Representative longitudinal flow-cytometry analysis of dissociated organoid cells (Bas, Basal; LumP, Luminal Proximal; PrU, Periurethral; Lum Distal, Luminal Distal). **d.** End-point RT-PCR analysis for selected marker genes. **e.** Bulk-RNAseq analysis (n=3; individual data points are shown with bar graphs representing mean value). **f.** Representative Haematoxylin-Eosin (H&E) staining of mouse prostate tissue and organoid sections (scale bars = 50 μm). **g.** Representative Immunofluorescence (IF) analysis for selected markers in mouse prostate tissue and organoid sections (scale bars = 50 μm). **h.** IF staining for selected markers in mouse prostate tissue and organoid sections (scale bars = 50 μm).

The epithelium of the prostate is characterized by a slow cellular turnover. In contrast, prostate organoids appeared to proliferate indefinitely – while retaining low levels of genomic instability (Supplementary Fig. S3) – raising the question of how culture conditions enable persistent cycling in a near-physiological manner. Either an excess of stimulatory cues in culture, or a lack of inhibitory ones – or both – may explain the shift from homeostatic quiescence *in vivo* to unrestrained mitotic activity *in vitro*. Using a ‘n −1 approach’ for assessing the requirement for growth factors and inhibitors in culture (Supplementary Fig. S4a), we found that prostate organoids are strictly dependent on Egf, with sub-nanogram concentration levels being sufficient for cell cycle progression (Supplementary Fig. S4b,c). Still – based on ligand/receptor expression patterns – Egf signalling alone is unlikely to explain the excess of proliferation in culture. Indeed, *Egf* is highly transcribed by distal luminal cells in the adult prostate^28^, and progenitor cells express the Egf receptor *in vivo* – independently of their proximal or distal location – at levels comparable to those observed *in vitro* (Supplementary Fig. S4d). Therefore, we focused on the requirement for the Tgf-β receptor inhibitor A83-01 for the continuous expansion of mouse prostate organoid cultures.

### Intra-epithelial non-canonical Activin A signalling is a key mediator of the Tgf-β induced cytostatic response in mouse prostate organoids

Upon A83-01 withdrawal, organoids displayed a marked reduction in EdU incorporation within 24 hours, demonstrating a cytostatic response in this model (Fig. 2a, Supplementary Fig. S5a). A83-01 is a potent inhibitor of three type-I Tgf-β family receptors, and two of them - Acvr1b and Tgfbr1 (also known as Alk4 and Alk5) – were found to be expressed in prostate organoids (Supplementary Fig. S5b). Paracrine and autocrine ligand-receptor interactions have been demonstrated to negatively regulate epithelial proliferation^16,39^ leading us to investigate the release of Tgf-β ligands in organoid cultures. We employed a click-chemistry approach to enrich for secreted proteins released in the culture medium, followed by mass-spectrometry analysis^40^. Over multiple experiments, we consistently recovered Activin A peptides (encoded by the *Inhba* gene) in the organoid supernatant, and only in one instance, a Tgfb1 peptide (Fig. 2b). We were also able to immunolocalize Acvr1b in organoid cells, and in progenitor cells *in vivo* (Fig. 2c), with an expression pattern similar to Egfr (Supplementary Fig. S4d). Replacement of A83-01 with Follistatin - a well-known Activin A inhibitor^41^ – was sufficient to sustain proliferation, revealing a prominent role for the latter in inducing a cytostatic response (Fig. 2d). To gain greater insights on the downstream pathway, we boosted Tgf-β receptor activity by combining A83-01 withdrawal with the supplementation of distinct ligands. Activin A was found to enhance the non-canonical arm of Tgf-β signalling – mediated by the Tgf-β activated kinase, Tak1 (encoded by the *Map3k7* gene), and the downstream p38 MAPKs – as well as the accumulation of the cell cycle inhibitor p21 (Fig. 2e). In contrast, Tgfb1 increased the activity of the canonical Tgf-β pathway – via Smad2/3 phosphorylation, but with little, if any, alteration of p21 levels. This dichotomy prompted us to functionally test the role of the canonical, and non-canonical pathway, respectively. Disruption of the canonical pathway, via shRNA-mediated silencing of Smad4, did not alter the cytostatic response upon A83-01 withdrawal (Supplementary Fig. S6). In contrast, the inhibition of either Tak1 or the structurally related p38α and p38β MAPKs - using a variety of inhibitors (Fig. 2f,g; Supplementary Fig. S7a,b) - was sufficient to ensure organoid expansion in the absence of Tgf-β receptor blockade.

**Figure 2.**
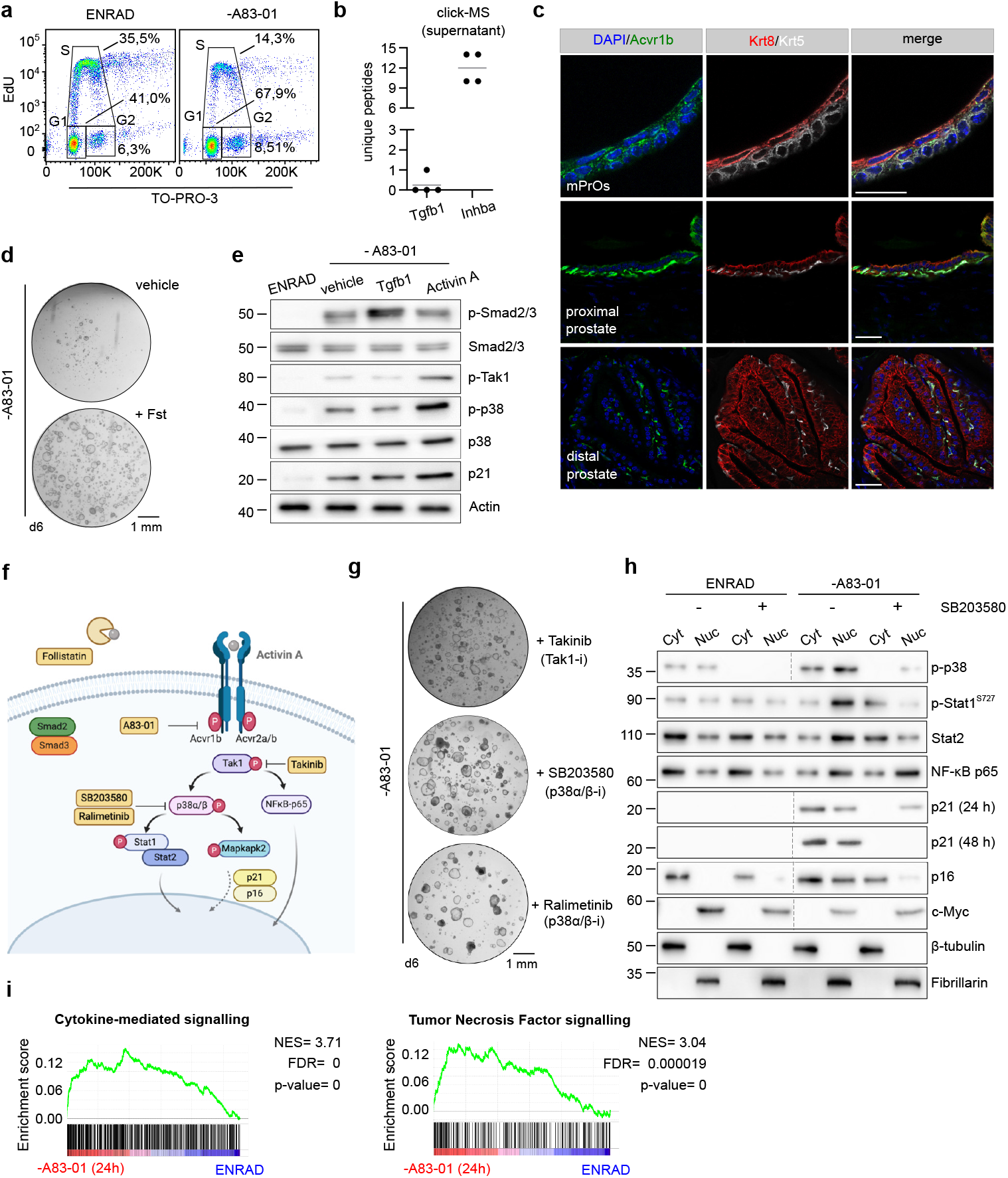
Intra-epithelial non-canonical Activin A signalling mediates the Tgf-β induced cytostatic response in mouse prostate organoids. **a.** Cell cycle analysis by flow cytometry (EdU *vs.* TO-PRO-3) in complete medium (ENRAD) or in the absence of A83-01 (24 hours). **b.** Detection of proteins secreted by mouse prostate organoids in culture based on click-chemistry enrichment and mass spectrometry analysis (n=4; data points are shown with crossing line representing mean value). **c.** IF analysis for selected markers in mouse prostate tissue and organoid sections (scale bar = 50 μm). **d.** Representative stereoscopic images of mouse prostate organoids cultured in the absence of A83-01, with or without Follistatin (Fst, 500 ng/mL, 6 days). **e**. Western blot (WB) analysis in mouse prostate organoids for selected canonical and non-canonical Tgf-β signalling mediators, and the cell cycle inhibitor p21 (Activin A, 50 ng/mL; Tgfb1, 500 ng/mL; 24 hours). **f.** Schematic view of the non-canonical Activin A pathway, including inhibitors used for the experiments described in this figure. **g**. Representative stereoscopic images of mouse prostate organoids following treatment with Takinib (Tak1 inhibitor; 5 μM, 6 days), SB203580 (p38α/β inhibitor; 10 μM, 6 days) or the structurally unrelated Ralimetinib (p38α/β inhibitor; 1 μM, 7 days). **h**. Nuclear fractionation and WB analysis in mouse prostate organoids for selected signalling mediators and cell cycle regulators, in the presence or absence of SB203580 (p38α/β inhibitor; 10 μM, 24 hours). **i**. Gene set enrichment analysis (GSEA) plots displaying significant enrichment for cytokine-mediated and Tumour Necrosis Factor (TNF) signalling in mouse prostate organoids cultured without A83-01 (24 hours) versus complete medium (ENRAD).

TGF-β receptors, and cytokine-stimulated receptors (*e.g.*, IL-1, TNF, type-I interferons), are known to converge on TAK1-p38 MAPK signalling to activate a variety of downstream factors controlling immune- and stress-related responses, including well-characterized transcriptional programmes^42–45^. By combining bulk-RNA sequencing and biochemical approaches, we confirmed that in the absence of Tgf-β receptors inhibition, Tak1-p38α/β signalling resulted in phosphorylation and nuclear shuttling of immune-related transcription factors (*e.g*., Stat1/2, NF-kB), which led to the transcription of immune related gene sets (*e.g.*, induced by TNF or type-I interferons)^46^, as well as to the phosphorylation of the stress-related kinase Mapkapk2 (Fig. 2h,i; Supplementary Fig. S7c-f). Importantly, Tak1-p38 MAPK activity was associated with the nuclear accumulation of the key cell cycle inhibitors p21 and p16 (Fig. 2h).

Collectively, our data demonstrated that intra-epithelial non-canonical Activin A signalling induce a cytostatic response in mouse prostate organoids.

### Evasion from the Tgf-β induced cytostatic response via downregulation of intra-epithelial Activin A signalling

We reasoned that our biobank may offer the opportunity to discover mechanisms of evasion from the Tgf-β induced cytostatic response, mediated by intra-epithelial Activin A signalling, in an unbiased manner. Therefore, we attempted to culture multiple prostate organoid lines in the absence of A83-01, waiting for the potential emergence of clones capable to thrive in these conditions. Out of nine prostate organoid lines (from three distinct mice), six were irreversibly lost within few weeks. Still, three lines survived for an extended period, with two of them eventually adapting to the absence of A83-01 and recovering the ability to be passaged at clonal density (Fig. 3a,b; Supplementary Fig. S8). We performed bulk-RNA sequencing on the C57#1 DLP organoid line in the presence of A83-01, one day after inhibitor withdrawal, and upon adaptation; additionally, we sequenced the C57#3 DLP organoid line in control conditions and following adaptation (Fig. 3c). We focused on transcriptional alterations shared by both lines displaying adaptation to A83-01 withdrawal. First, we noticed that gene signatures associated with ATR signalling were strongly upregulated upon adaptation (Fig. 3d), indicating potential DNA replication stress in S-phase, a finding consistent with phosphorylation of the cell cycle Checkpoint kinase 1 (Chek1) (Fig. 3e). Of note, in the adapted C57#1 DLP line we detected widespread genomic instability and telomere doublets, a hallmark of DNA replication stress, while retaining an intact p53 pathway (Fig. 3f-h; Supplementary Fig. S9). Second, in both adapted organoid lines, we observed the downregulation of immune-related transcriptional programmes (*e.g.*, type-I interferon stimulated genes), suggesting an impairment of non-canonical Activin A signalling (Supplementary Fig. S10). Consistently, we found that adapted organoid lines significantly reduced Activin A secretion in culture (Fig. 3i), and that exogenous Activin A (but not Tgfb1) was sufficient to restore the cytostatic response (Fig. 3j,k; Supplementary Fig. S11).

**Figure 3.**
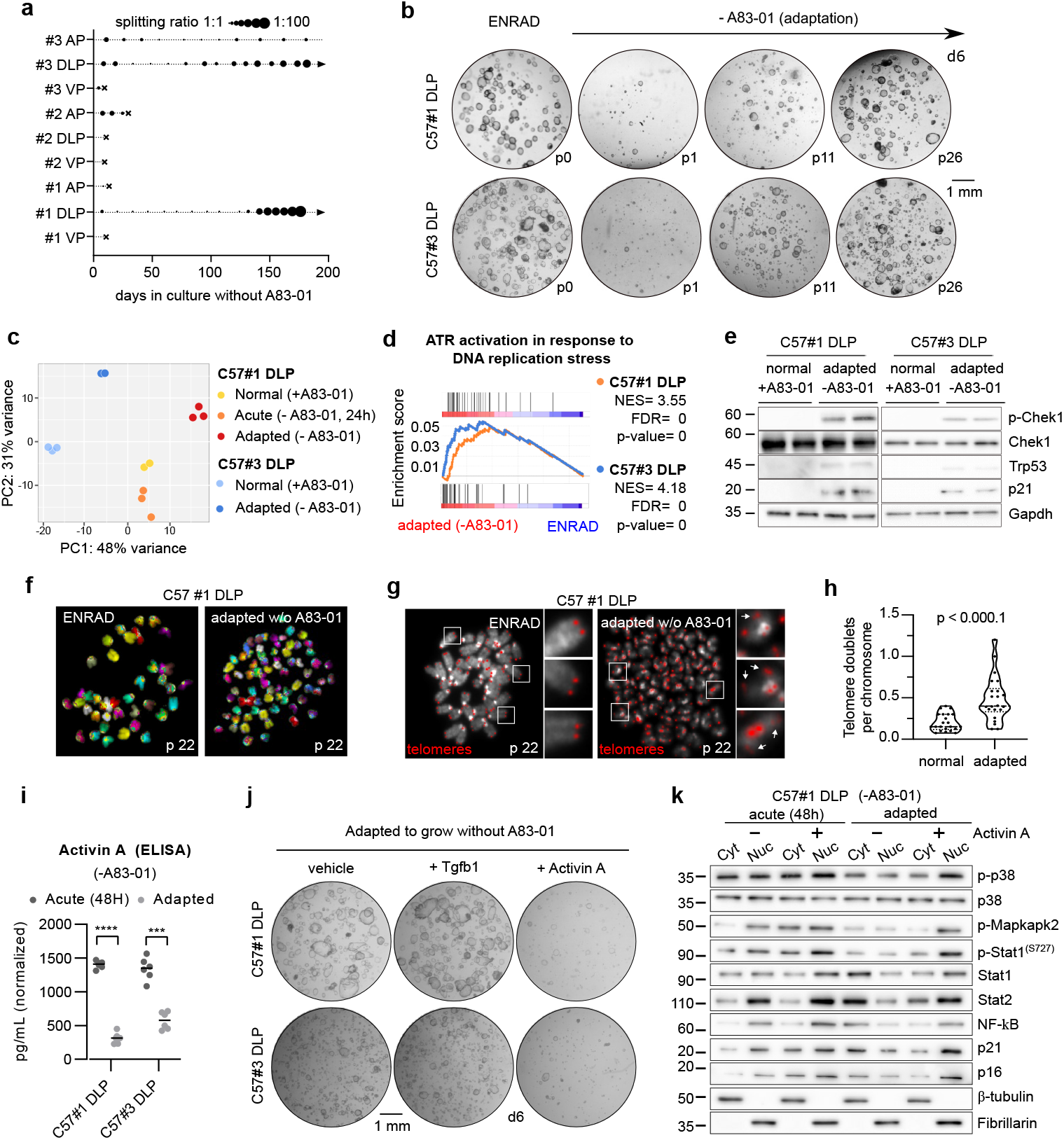
Evasion from the Tgf-β induced cytostatic response occurs via downregulation of intra-epithelial Activin A signalling and leads to DNA replication stress and genomic instability. **a.** Diagram depicting the expansion of mouse prostate organoid cultures in the absence of the Tgf-β ligand inhibitor A83-01 (arrow = continuous expansion; dot = passage; cross = culture loss). **b.** Representative stereoscopic images of C57BL/6J DLP #1 and # 3 mouse prostate organoid lines during adaptation in the absence of A83-01. **c.** Principal Component Analysis (PCA) based on bulk RNA-sequencing of C57#1 DLP and C57#3 DLP mouse prostate organoids grown in normal conditions (ENRAD), upon acute A83-01 depletion (-A83-01, 24 hours) (C57#1 DLP only) or following adaptation (-A83-01, long-term). **d.** Gene set enrichment analysis (GSEA) plot displaying significant enrichment for activation of ATR signalling in mouse prostate organoid lines (C57#1 DLP, top; C57#3 DLP, bottom) adapted to grow in the absence of A83-01 *vs*. normal control organoids cultured in complete medium (ENRAD). **e.** WB analysis in C57#1 and C57# 3 DLP mouse prostate organoid lines adapted to grow in the absence of A83-01 *vs.* normal control organoids cultured in complete medium (ENRAD). Immunoblots are displayed for Chek1 (ATR signalling mediator), Trp53 and p21. **f.** Representative spectral karyotype (SKY) images of metaphase spreads obtained from C57#1 DLP mouse prostate organoids cultured in normal conditions (ENRAD) or upon adaptation without A83-01. Widespread genomic instability is observed following adaptation. **g-h.** Representative telomere FISH images - and quantification - in C57#1 DLP mouse prostate organoids cultured in normal conditions (ENRAD) or upon adaptation without A83-01. Widespread telomeric instability is observed following adaptation. **i.** Enzyme-linked immunosorbent assay (ELISA) for Activin A expression in the supernatant of C57#1 and C57# 3 DLP mouse prostate organoid lines, upon acute A83-01 removal (48 hours) or long-term adaptation. Two-way ANOVA, Sidak’s test, p-value *** (<0.001), **** (<0.0001). **j.** Representative stereoscopic images of C57#1 and C57# 3 DLP mouse prostate organoid lines adapted to the absence of A83-01 and subsequently treated with either Tgfb1 (500 ng/mL, 6 days) or Activin A (50 ng/mL, 6 days). **k.** WB analysis in the C57#1 DLP mouse prostate organoid line upon acute removal (24 hours) or long-term adaptation to A83-01 removal, in the presence or absence of Activin A (50 ng/mL, 24 hours).

Thus, mouse prostate organoids are capable to dampen the Tgf-β induced cytostatic response by downregulation of intra-epithelial Activin A signalling.

### Mouse prostate organoids with reduced intra-epithelial Activin A signalling display enhanced engraftment upon syngeneic transplantation

We wondered whether a reduced intra-epithelial Activin A signalling may release the progenitor proliferative potential in response to basal growth stimuli, within the relative quiescent microenvironment of the adult prostate epithelium. To test this hypothesis, we orthotopically transplanted dissociated mouse prostate organoid cells into immunocompetent syngeneic mice (Fig. 4a). Donor cells were injected into the extensive branchial structures of the distal anterior prostate, to avoid damage to the delicate proximal ducts, and maximize the probability of retention. Implantation of donor cells into an immunocompetent host tissue characterized by a slow turnover can be considered challenging. Still, we found that, in comparison to control organoid cells, organoid cells adapted to grow without A83-01 implanted with high frequency (Fig. 4b), and gave rise to dysplastic foci, characterized by elevated mitotic index, cuboidal histology and nuclear atypia (Fig. 4c,d).

**Figure 4.**
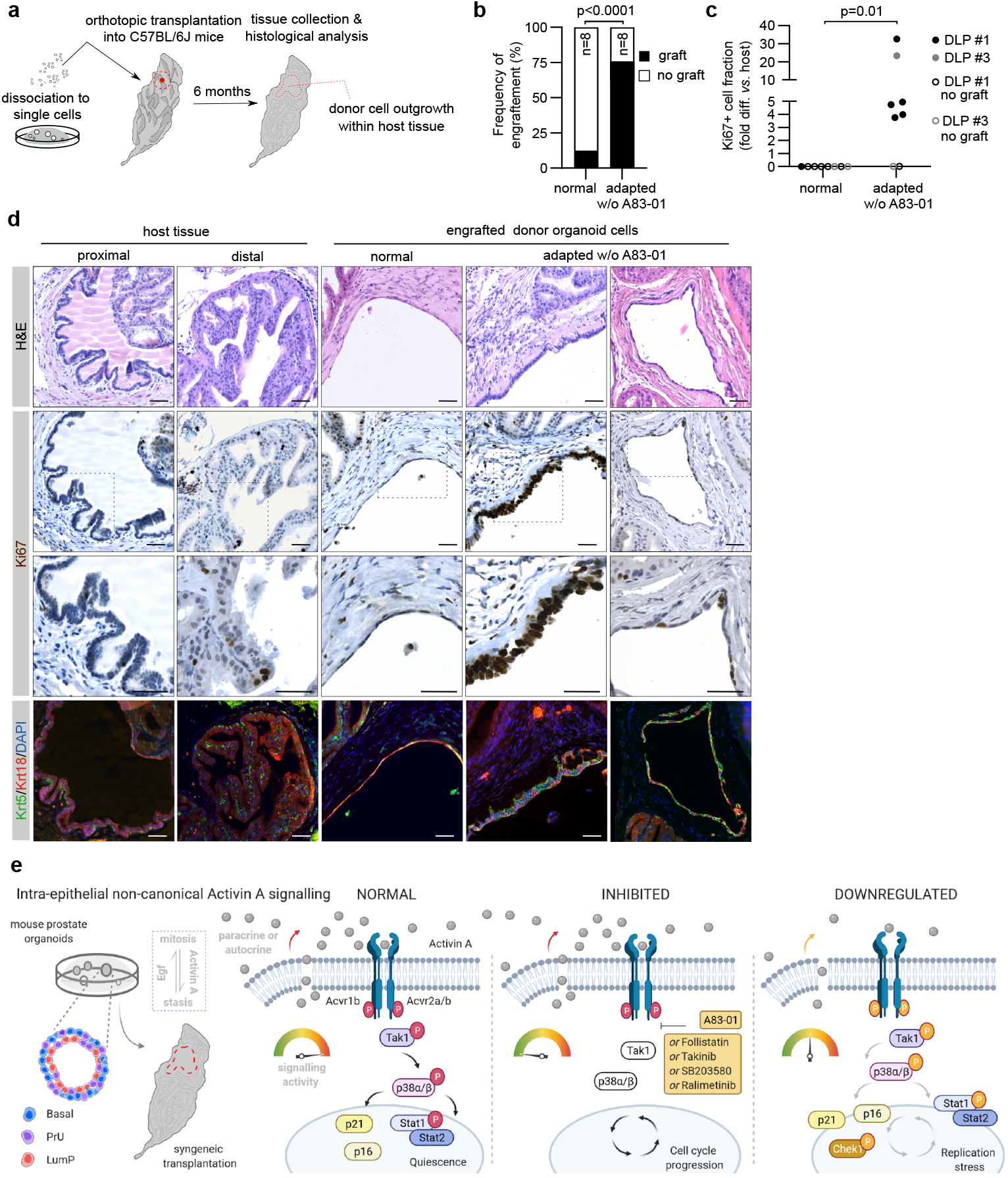
Enhanced engraftment of mouse prostate organoids with reduced Activin A signalling into immunocompetent hosts. **a.** Schematic view of the orthotopic transplantation strategy into the anterior prostate lobe of immunocompetent syngeneic C57BL/6J mice. **b.** Frequency of engraftment (%) of mouse prostate organoid lines (C57#1 DLP and C57#3 DLP) expanded in complete medium (normal) or adapted to the absence of A83-01 (binomial test (two-tailed)). **c.** Normalized mitotic (Ki67+) cell index in grafts of mouse prostate organoid lines (C57#1 DLP and C57# 3 DLP) expanded in complete medium conditions (normal) or adapted to the absence of A83-01 (non-parametric Mann-Whitney test). **d.** Representative H&E, IF, and immunohistochemistry (IHC) analyses of host prostate tissue and engrafted donor organoid cells (scale bar = 50 μm). **e.** Intra-epithelial non-canonical Activin A signalling safeguards prostate progenitor quiescence. In normal circumstances, autocrine or paracrine Activin A triggers Tak1/p38 Mapk non-canonical signalling - antagonizing the pro-proliferative Egf pathway and enforcing cellular quiescence. Mechanistically, Tak1/p38 Mapk activity leads to p16 and p21 nuclear translocation - as well as to Stat1/2-dependent transcription of type-I interferon genes. Prostate organoid cultures require the suppression of this non-canonical signalling for continuous expansion, which can be achieved at different levels of the signalling cascade, using multiple inhibitors. Organoids with reduced levels on intra-epithelial Activin A signalling can be selected *in vitro*, and display enhanced engraftment efficiency *in vivo*, upon orthotopic transplantation into syngeneic hosts. Notably, the partial down-regulation of this pathway is associated with DNA replication stress and genomic instability.

We conclude that intra-epithelial non-canonical Activin A signalling safeguards quiescence in prostate progenitors.

## Discussion

Signalling pathways ensure coordination of tissue development, homeostasis, regeneration, and their disruption can lead to disease. The molecular bases of specific signals are difficult to investigate, due to the challenges of disentangling cellular cross talks *in vivo*, and of establishing representative models *in vitro*. More recently, organoid models in defined media conditions have opened new opportunities for the study of epithelia^47^. Benchmarking of these models with their corresponding *in vivo* counterpart is paramount for the correct experimental interpretations^48^. Here, we demonstrated that mouse prostate organoid cultures enable the continuous expansion of epithelial progenitors *in vitro*, including basal, LumP and PrU cells. Such cell types are predominantly found near the urethra *in vivo*, but also in the distal prostate compartment at low frequency. Our work revealed that progenitor proliferation is dynamically regulated by the antagonistic equilibrium between Egf and non-canonical Activin A signalling, respectively – with at least partial reduction of the latter required for cell cycle progression. The rationale perturbation of additional biochemical signals, and mechanical cues, may enhance progenitor differentiation towards distal luminal cells in culture.

It has long been known that the broad family of TGF-β signals induces a cytostatic response in a large variety of epithelial cells, but the specific pathway acting in the prostate has remained poorly understood. Earlier studies pointed to the importance of the Tgf-β family ligand Activin^49,50^. At the mechanistic level, DePinho and colleagues initially focused on the role of Smad4 as a proliferative barrier in a *Pten-loss* driven mouse model of prostate cancer^51^. Follow-up studies from the same group and others, carried out in humans and mice, have led to a more complex view^52,53^, with the involvement of both canonical and non-canonical pathways.

We propose a prominent role for non-canonical Activin A signalling in safeguarding quiescence in prostate epithelial progenitors (Fig. 4e). Our model may be relevant beyond tissue homeostasis and the response to inflammation. Genes encoding for core components of the non-canonical Activin A signalling pathway (*e.g*., *ACVR2A, MAP3K7*) are frequently lost in prostate cancer, based on large cohorts of patients in the U.S.A.^54^ and in China^55^. Moreover, *MAP3K7* loss has been linked to genomic instability in human prostate cancer cell lines^56^ and found to promote an aggressive transcriptional programme in prostate tumours, based on a recent systematic pan-cancer analysis^57^. In contrast, genetic alterations rarely affect TGFB1 receptors (*e.g*., *TGFBR2*) or SMAD factors (*e.g., SMAD4*), and enhanced canonical TGF-β signalling has been reported in metastatic biopsies in therapy-resistant prostate cancer patients^58^. We speculate that loss of the non-canonical TGF-β arm could impair the cytostatic response, while sparing the well-known transforming potential of the TGF-β canonical pathway.

Tak1/p38-MAPK signalling stimulated two main sets of effector proteins in prostate progenitors. On the one hand, we observed the nuclear translocation of key negative cell cycle regulators (*e.g*., p16 and p21). On the other hand, using biochemical and transcriptional analyses, we demonstrated the activation of a broad transcriptional response, reminiscent of those induced by inflammatory cytokines and pathogens. These findings are in line with recent observations on the immune function of structural cells^59^ and suggest a cross talk between the epithelial and immune compartments, beyond the well-known mechanisms of anti-microbial defence. Prostate progenitors – and, perhaps, other types of epithelia cells – may have the ability to signal changes in their proliferative status and, immune cells may have the capability to adjust their function in response.

To test the relevance of intra-epithelial non-canonical Activin A signalling for the enforcement of quiescence in prostate progenitors, we performed experiments *in vitro* and *in vivo*. Our long-term organoid cultures – in the absence of Tgf-β receptor blockade – revealed that cells capable to re-enter cell cycle had downregulated Activin A secretion. Moreover, those cells were also capable to implant and proliferate at increased frequency *in vivo*. While stromal sources of Tgf-β ligands have been previously described in the prostate^25,26^, our study is the first to demonstrate a key role for intra-epithelial signalling.

Notably, dysregulation of non-canonical Activin A signalling was associated to DNA replication stress and genomic instability, a finding that may be relevant for tumour initiation. Indeed, distal LumP cells have been shown to serve as cell-of-origin for prostate cancer^30^. In this regard, our orthotopic transplantation approach may be particularly relevant for investigating the tumorigenic potential of distal progenitor cells.

Finally, P38 MAPK inhibitors – including Ralimetinib – are currently being tested in phase 1/2 clinical trials^60^. Quiescent tumour progenitors – induced by the broad family of TGF-β signals - are emerging as key mediators of chemotherapy resistance in solid malignancy^61^. In advanced prostate cancers with a genetically intact Activin A non-canonical pathway, P38 MAPK inhibitors may force tumours progenitors out of quiescence, improving the efficacy of standard chemotherapy regimens^62^. While the complexity and pleiotropy of TGF-β signalling has historically complicated drug development^63^, the elucidation of cell- and context-specific pathways may lead to novel therapeutic opportunities.

## Methods

### Mice

Animal experiments were performed according to the European Communities Council Directive (2010/63/EU) and approved by the Italian Ministry of Health and the University of Trento Animal Welfare Committee (642/2017-PR). Wild-type C57BL/6J (JAX # 000664) mice were purchased from the Jackson Laboratory. Wild-type BALB/c (CRL # 028) and CD-1 (CRL # 022) mouse strains were obtained from the Charles River Laboratories.

### Isolation of mouse prostate tissue

The anterior (AP), dorso-lateral (DLP) and ventral prostate (VP) lobes were dissected individually, using a transverse cut at the intersection of each lobe with the urethra. Paired lobes were collected for organoid cultures, histology and immunostaining studies.

### Dissociation of mouse prostate tissue to single cells

Prostate tissue was minced into small pieces, washed, resuspended into a digestion buffer-including Collagenase II (1 mg/mL; Life Tech, 17101015) and Dispase II (10 mg/mL; Life Tech 17105041), and transferred into a gentleMACS C tube (Miltenyi Biotec). Tissue fragments were processed by alternating mechanical disruption - using the gentleMACS Dissociator (A.01-C tube programme) – and enzymatic digestion-incubating the solution at 37 °C on a tube rotator for 15 minutes. After three cycles, the cell suspension was pelleted, resuspended in TrypLE (Life Tech, 12605010), and incubated for 5 minutes at 37 °C. After two washes, the cell suspension was filtered through a 70 μm strainer and counted.

### 3D prostate organoid cultures

Dissociated prostate cells were resuspended in 80% growth factor-reduced basement matrix (either Matrigel^®^,Corning, 356231; or BME-2^®^,AMSBIO, 3533) and seeded at the concentration of approximately 50,000 cells/mL, by depositing at least six 40 μL drops at the bottom of a non-tissue culture treated plate. Basement matrix domes were left to solidify for 15 minutes and covered with ENRAD medium – including Egf (50 ng/mL; PeproTech, 315-09), Noggin (100 ng/mL; PeproTech, 120-10C), R-Spondin1 (10% conditioned medium), A83-01 (200 nM; Tocris, 2393) and dihydrotestosterone (10 nM; Merck, 10300) – supplemented with Y-27632 (10 μM; Calbiochem, 146986-50-7), as previously described^64^. Organoids were cultured in a standard tissue culture incubator, with medium replacement every 2-3 days. After 6 days from the initial seeding, organoids were imaged with a Leica MZ16F stereomicroscope and organoid forming efficiency was calculated. For subsequent passages, the basement membrane was dissolved using a recovery solution – including Dispase II (1 mg/mL) – and organoids were dissociated to small clumps/single cells as described above, using TrypLE. Following the first passage, organoids were seeded at the concentration of approximately 25,000 cells/mL.

### Lentiviral transduction of organoids

Organoids were dissociated to single cells, and approximately 50,000 cells were transduced for each condition. Spinoculation was performed in a low-adhesion 96 well-plate using 0.6 RTU of lentiviral solution, supplemented with polybrene (4 μg/mL; Sigma Aldrich, H9268) and complete medium (ENRAD) to reach a final reaction volume of 300 μL. The plate was sealed with parafilm and centrifuged for 1 hour at 600 g. Afterwards, the cells were resuspended in 200 μL of complete medium (ENRAD), supplemented with Y-27632 (10 μM), and incubated in suspension at 37 °C for 4-6 hour. After centrifugation, the cell pellet was resuspended in 80% basement matrix and seeded as described above. Antibiotic selection was initiated two days post-transduction. The following plasmids were used: pLenti-AIB-EGFP (kindly donated by Massimo Pizzato), pSUPER-retro-puro-Smad4 (Addgene #89829) and pSUPER-retro-puro-GFP shRNA (Addgene #30519).

### Treatments with growth factors and small molecule inhibitors

Growth factors and small molecules used in this study are described in Supplementary Table 1.

### Flow cytometry

Organoids were dissociated to single cells as described above. For cell surface antigen expression analysis, cells were incubated with anti-Cd24a-APC and Sca-1-PE-Cy7 antibodies (1:800 dilution) at 4°C for 20-30 minutes, followed by one wash with FACS buffer (1% FBS, 1mM EDTA). Cells were resuspended in FACS buffer, supplemented with 1 μM propidium iodide (Life Tech, P3566) for dead cell exclusion, before proceeding to the analysis. For DNA content analysis, cells were resuspended in 100 μl ice-cold PBS and transferred to a 15 ml tube. About 900 μl of ice-cold EtOH 70% were added dropwise while agitating the cell suspension on a vortex. Cells were then fixed for at least 2 hours at −20°C before proceeding with 3 washing steps in PBS, alternated by centrifugation (700 g, 5 min) with no brake. Afterwards, cell pellet was resuspended in 100 μL of DNAse-free RNAseA (0.5 μg/mL; Life Tech, 12091021) and incubated for 10 min at 37 °C. Cells were incubated with 100 μL of propidium iodide (50 μg/mL) for 30 minutes, at room temperature, before proceeding to the analysis. For cell cycle analysis, organoids were treated with 10 μM EdU for 3 hours. Afterwards, organoids were harvested, dissociated into single cells, and filtered through a 30 μm strainer. Cells were pelleted and stained with the Click-iT™ Plus EdU Alexa Fluor™ 488 Flow Cytometry Assay Kit (Thermo Fisher Scientific, C10632), according to the manufacturer protocol. After the incorporation of the fluorescent probe, cells were incubated with TO-PRO™-3 Iodide (Life Tech, T3695) to stain for DNA content, before proceeding to the analysis. Flow cytometry was performed with a FACS Canto (BD) analyser, and data analysed with Flow Jo v.10.

### Histology, immunostaining, and live imaging

Organoids were cultured for 5-7 days, released from the basement membrane as described above, seeded in a neutralized collagen type-I solution (Corning, 354249), and cultured for additional 24 hours, before proceeding to fixation in 4% paraformaldehyde (Sigma Aldrich, P6148) for 5 hours, at room temperature. Prostate tissue was harvested and immediately fixed using the same conditions. Paraffin embedding and 5 μm sectioning were carried out according to standard procedures. For immunofluorescence studies, antigen retrieval was performed using a citrate-based buffer (pH 6.0) (Vector Lab, H3300). Slides were incubated in blocking solution (5% FBS + 0.1% Triton-X in PBS), before proceeding to staining with primary antibodies, at 4°C, overnight. After three washes, spectrally distinct fluorochrome-conjugated antibodies were incubated for 2 hours at room temperature. After three additional washes, samples were counterstained with Hoechst 33342 (Abcam, ab145597), and the coverslip was applied, using FluorSave mounting medium (Merck, 345789). For immunohistochemistry studies, a similar protocol was followed, but using biotin-conjugated secondary antibodies. The detection was performed using the Vectastain^®^ Elite ABC Peroxidase kit (Vector Labs, PK-6100) according to the manufacturer instructions. The final reaction was blocked by washing slides with water, and coverslips were applied using the DPX mounting medium (Sigma, 06522). For haematoxylin and eosin (H&E) staining, deparaffinised sections were incubated with Gill haematoxylin (Merck, GH5232) for 2 minutes and washed with water. Samples were washed with ethanol, incubated with eosin Y (Merck, HT110132) for 3 minutes, washed again twice with ethanol, and treated with xylene, before mounting the coverslips in phenol based mounting medium. For immunostaining with anti-Egfr and anti-Acvr1b antibodies, the urogenital apparatus was isolated, snap-frozen in 2-methyl-buthanol cooled in liquid nitrogen, and cryo-sectioned at 20 μm. Tissue slides were fixed in 4% paraformaldehyde for 20 minutes at room temperature, before proceeding as described above. For live imaging, organoids were stably transduced with pLenti-AIB-EGFP. Images were acquired using either a Zeiss Axio Imager M2, or a Zeiss Axio Observer Z1 Apotome, or a Leica TCS SP8 Confocal. Image analysis was performed with the Zeiss ZEN software or ImageJ (v.2.0.0-rc-69/1.52i)^65^. Primary antibodies are listed in Supplementary Table 2.

### RNA extraction

Total RNA was extracted using the RNeasy Plus Micro kit (Qiagen, 74034) according to the manufacturer instructions, and analysed with an Agilent BioAnalyzer 2100 to confirm integrity (RIN > 8), before proceeding with downstream applications.

### End-point semi-quantitative and quantitative real-time PCR

RNA was retrotranscribed into cDNA using the iScript™ cDNA synthesis kit (BioRad, 1708891). End-point PCR was performed using Phusion Universal qPCR Kit (Life Tech, F566L), with PCR products visualized by standard gel electrophoresis. For quantitative real-time gene expression analysis, the qPCRBIO SyGreen Mix (PCRBiosystems, PB20.14-05) was used according to the manufacturer instructions. At least three independent biological replicates were run for each sample, using the CFX96 Real Time PCR thermocycler (Bio-Rad). The data were processed using Bio-Rad CFX Manager software (v.3.1), while gene expression quantification and statistical analyses were performed with GraphPad PRISM (v.6.01). Primer sequences are included in Supplementary Table 3.

### RNA sequencing and data analysis

cDNA libraries were prepared with TruSeq stranded mRNA library prep Kit (Illumina, RS-122-2101) using 1 μg of total RNA. RNA sequencing was performed on an Illumina HiSeq 2500 Sequencer using standard Rapid Run conditions at the Next-Generation Sequence Facility of University of Trento. The obtained reads were 100 bp long, single ends, and 25 million on average for each sample. FASTQ file from Illumina HiSeq2500 sequencing machine underwent adapter removal and quality-base trimming using Trimmomatic-v0.35. Genomic alignments were performed onto the Mouse genome (mm10 assembly version) using STAR-v2.6.0 aligner with a maximum mismatch of two and default settings for all other parameters. Then, uniquely mapped reads were selected and processed with HTSeq-count v0.5.4 tool to obtain gene-level raw counts based on GRCm38.92 Ensembl (www.ensembl.org) annotation. Genes with CPM (Counts Per Million) < 1 in all replicates were considered unexpressed and hence removed from the analysis. TMM (Trimmed Mean of M values) normalization and CPM conversion were next performed to obtain gene expression levels for downstream analyses. For each comparison, differential expression testing was performed using the edgeR-3.20.9 statistical package. According to the edgeR workflow, both common (all genes in all samples) and separate (gene-wise) dispersions were estimated and integrated into a Negative Binomial generalized linear model to moderate gene variability across samples. For each comparison, genes having a log Fold-change outside the range of +/-1.5 and a FDR q-value equal or smaller than 0.01 were considered as differentially expressed between the two groups.

### Gene Set Enrichment Analysis (GSEA)

For the gene set enrichment analysis, the GSEA software (v4.0.3) was run in the ‘pre-ranked’ mode using the Fold-change as a ranking metric and an FDR enrichment threshold of 0.25. Gene sets were directly obtained from the Molecular Signature (MSig) database (http://software.broadinstitute.org/gsea/msigdb) focusing on all available sets reported in the following MSigDB collections: C2 (curated gene sets): Biocarta, Kegg, Reactome; C5 (Gene ontology): Biological Processes, Cellular Component, Molecular Function; C6 (oncogenic signatures) and C7 (immunologic signatures).

### Principal Component Analysis (PCA)

PCA was performed using the DESeq2 R-package^66^ as follows: normalized counts (CPM) were firstly converted into a DESeqDataset object through a DESeqDatasetFromMatrix function with default parameters and transformed through the variantStabilizingTransformation function to stabilize variance-mean relation across samples. Then, transformed data was analyzed by PCA (plotPCA function) generating a two-dimensional space where the two first components are represented.

### Subcellular Fractionation and Western blotting

Cell pellets from organoid cultures were obtained as previously described and lysed in fresh RIPA buffer (50 mM Tris-HCl, pH 7.5, 150 mM NaCl, 1% Triton X-100, 1% sodium deoxycholate, 1% NP-40) supplemented with protease (Halt™ protease inhibitor cocktail, Life Tech, 87786) and phosphatase inhibitors (Phosphatase-Inhibitor Mix II solution, Serva, 3905501). Nuclear/cytoplasmic fractionation was performed using NE-PER Nuclear and Cytoplasmic Extraction Kit (Life Tech, 78833) according to the manufacturer instructions. Protein concentrations were measured using the BCA Protein Assay Kit (Pierce™ BCA Protein Assay kit, Thermo Fisher Scientific, 23225) and a Tecan Infinite M200 Plate Reader.

Protein extracts were resolved via SDS-PAGE, transferred to polyvinylidene difluoride (PVDF) membrane (Merck, GE10600023) using a wet electroblotting system (Bio-Rad). The membranes were blocked with 5% non-fat dry milk or 5% BSA in TBS-T (50 mM Tris-HCl, pH 7.5, 150 mM NaCl, 0.1% Tween20) for 1 hour, at room temperature, and then incubated with gentle shaking with designated primary antibodies overnight, at 4°C. Membranes were incubated with HRP-conjugated secondary antibody in blocking buffer for 1 hour at room temperature. Immunoreactive bands were detected using ECL LiteAblot plus kit A+B (Euroclone, GEHRPN2235) with an Alliance LD2 device and software (UVITEC). Primary antibodies are provided in Supplementary Table 4.

### Click-it chemistry-based mass spectrometry analysis

Organoids were seeded at the approximate concentration of 50,000 cells/ml, depositing seven 40 ul domes *per* individual well of a 6-well non-tissue culture plate. Three wells were used for each condition. Following methionine depletion (2 hours), organoids were grown overnight at 37 °C with L-azidohomoalanine (AHA) medium. Conditioned supernatants were collected, supplemented with protease inhibitors, and stored at −80 °C until further processing. CLICK-IT enrichment of AHA-labelled secreted proteins was performed with the Click-iT™ protein enrichment kit (Thermo Fisher Scientific, C10416) as previously described^67,68^. Following trypsin digestion, peptides were purified by reversed-phase (C18) stage-tip purification^69^. LC-MS/MS analysis was performed by an EASY-LC 1000 coupled to a Q-Exactive mass spectrometer (Thermo Fisher Scientific). LC-MS/MS data analysis was conducted using the MaxQuant/Perseus software suite.

### Enzyme-linked immunosorbent assay (ELISA)

Activin A quantification was performed using the corresponding Quantikine ELISA Kit (R&D Systems, DAC00B) according to the manufacturer instructions.

### Karyotype analysis

Organoid cultures were treated with nocodazole (15 μM; Sigma, SML1665) for 5 hours. Organoids were recovered from basement membrane, and 900 μL/sample of 50 mM KCl were added to the cell pellet dropwise, followed by 10 minutes of incubation at 37 °C. After centrifugation (200 g, 5 minutes) 900 μL/sample of Carnoy’s fixative (methanol/acetic acid 3:1) was added dropwise. Samples were resuspended and incubated for 10 minutes at 37 °C, followed by three washes with methanol/acetic acid 2:1. Approximately 25,000 cells/samples were resuspended in 50 μL and dropped from at least 1 meter of height, directly on a glass slide. After air-drying, the glass slide was incubated with Hoechst 33342, at room temperature, for 10 minutes, and then washed with methanol/acetic acid 2:1 for 5 minutes. After air-drying, coverslips were mounted with ProLong Gold Antifade (Invitrogen, P36934). Images were acquired at the Zeiss Observer Z1 microscope and analysed with ImageJ (v2.0.0-rc-69/1.52i)^65^.

### Multicolor FISH (M-FISH), Chromosome painting and Telomeric FISH

For M-FISH, fixed cells were dropped onto glass slides and hybridized with the 21XMouse Multicolor FISH Probe Kit (MetaSystems, D-0425-060-DI), as previously described^70^. Briefly, the slides were denatured in 0.07 N NaOH and then rinsed in a graded ethanol series. The probe mix was denatured using a MJ mini personal thermal cycler (Bio-Rad) with the following program: 5 minutes at 75 °C, 30 seconds at 10 °C, and 30 minutes at 37 °C. The probe was added to the slides and the coverslip was sealed using rubber cement. The samples were then hybridized in a humidified chamber at 37 °C for 48 h, washed in saline-sodium citrate (SSC) buffer for 5 min at 75 °C, and finally counterstained with DAPI (Abcam, 6843.2), in Vectashield mounting medium. Metaphases were visualized and captured using a Zeiss Axio-Imager M1 microscope. The karyotyping and cytogenetic analysis of each single chromosome was performed using the M-FISH module of the ISIS software (MetaSystems). A total of 25 metaphases for each sample spreads were analysed in two independent experiments.

For chromosome painting, fixed cells were dropped onto glass slides and hybridized with enumeration XMP painting probes specific for chromosomes X (red label) and chromosome Y (green label) (MetaSystems, D-1420-050-OR, D-1421-050-FI) following the manufacturer instructions. Briefly, probes were applied to the slides, denatured at 75 °C for 2 minutes, and then incubated at 37 °C overnight. The slides were washed in SSC and counterstained with DAPI in antifade reagent (MetaSystems, D-0902-500-DA). Metaphases were visualized and captured using a Zeiss Axio-Imager M1 microscope. A total of 100 metaphases were analysed for each sample in two independent experiments.

For telomeric FISH, staining was performed as previously described^71^. Briefly, slides and the Cy3 linked telomeric (TTAGGG)3 PNA probe, (DAKO Cytomatation, K5326) were co-denatured at 80 °C for 3 minutes, and hybridized for 2 hours at room temperature, in a humidified chamber. After hybridization, slides were washed and then dehydrated with an ethanol series and air dried. Finally, slides were counterstained with DAPI and Vectashield. Images were captured at 63× magnification using a Zeiss Axio-Imager M1 microscope, and the telomere signals were analysed using the ISIS software (MetaSystems). Telomere doublets frequency was calculated as the ratio between the number of doublets signals and the total number of chromosomes in each metaphase analysed^72^. At least 20 metaphases in two independent experiments were analysed.

### Orthotopic organoid transplantation

Orthotopic transplantation of organoids into the prostate of syngeneic immune-competent C57BL/6J mice was performed adapting a previously published method^73^. Organoids were dissociated as described above, with 50,000 cells *per* injection resuspended in 10 μL of 50% basement matrix, supplemented with methylene blue (as tracer). Upon abdominal incision of the host, the left anterior prostate lobe was exposed, and injected into the distal part. The contralateral lobe was injected with saline as negative control. Mice were regularly monitored and sacrificed after 6 months for tissue collection and histopathological analysis.

### Statistical analysis and reproducibility

No statistical methods were used to predetermine sample size. The *in vitro* experiments were not randomized, and the investigators were not blinded to allocation during experiments and outcome assessment. The *in vivo* transplantation experiments were randomized, and the investigators were blinded to allocation during experiments and outcome assessment. The *in vitro* experiments were carried out on organoid lines derived from at least two distinct animals and repeated at least three independent times. The *in vivo* transplantation experiments were based on two distinct organoid lines and were repeated at least two independent times. Data collection was performed using Microsoft Office Excel 2016–2018 and statistical analysis was performed using GraphPad Prism 6 software. The number of replicates, the format of the data, and the statistical tests are indicated in figure legends. p-values < 0.05 were considered significant.

## Data availability

RNA sequencing datasets have been deposited on BioProject with the dataset identifier PRJNA659468. All other data supporting the findings of this study are available from the corresponding authors upon reasonable request.

## Acknowledgments

We are grateful to Francesca Demichelis, Alberto Inga, Giannino del Sal, Marco Marcia, and Karuna Ganesh, for critical reading of the manuscript, and, Luciano Conti, Alessio Zippo, Luca Fava, Fulvio Chiacchiera, Luca Tiberi, Massimo Lopes, Alberto Briganti, Matteo Bellone and Marianna Kruithof-de Julio for fruitful discussions. We thank Francesca Demichelis, Massimo Pizzato, Luca Fava, Yari Ciribilli, and Valeria Poli, for sharing reagents. We thank Veronica de Sanctis, Roberto Bertorelli and Paola Fassan of the Next Generation Sequencing Facility of the University of Trento for RNA sequencing. We also thank Sergio Robbiati and Marta Tarter of the CIBIO Model Organism Facility, Marina Cardano of the CIBIO Cell Technology Facility, Giorgina Scarduelli and Michela Roccuzzo of the CIBIO Advanced Imaging Facility, Isabella Pesce of the CIBIO Cell Analysis and Separation Facility, Valentina Adami and Michael Pancher of the CIBIO High-Throughput Screening Facility, and Isabelle Bonomo for assistance with experiments. We are grateful to Qingwen Jiang and Ahmed Mahmoud for help with illustrations, which were created using Inkscape and Biorender.com. This study was primarily supported by grants from the Giovanni Armenise-Harvard Foundation (Career Development Award to A.L.), the Lega Italiana Lotta ai Tumori (LILT-Bolzano to A.L.), the Italian Ministry for Research and the University (MIUR PRIN 2017 to A.L.), the Associazione Italiana per la Ricerca sul Cancro (AIRC MFAG 2017-ID 206221 to A.R.), the NIH (R01CA238005 to M.M.S.) – and by core funding from the University of Trento. Individual fellowships were awarded from the United States Department of Defense (W81XWH-18-1-0424 to F.C.), the European Union (H2020-MSCA 749795 to A.Be.), the Fondazione Umberto Veronesi (FUV 2016 and 2017 to F.C., FUV 2016 to A.A., and FUV 2018 to A.Be.), and the University of Trento (Ph.D. fellowship to V.F, and D.D.F.).

## Author contributions

F.C. made initial observations and designed the project in consultation with A.L.; F.C., V.F. and A.L. designed the experiments with the contribution of A.Ala. and D.D.F. regarding the Tak1/p38 MAPK signalling; F.C., with the contribution of M.Z., generated and characterized the normal prostate organoid lines described in this study; F.C., with the contribution of M.D.P., carried out the wet-lab based experiments defining the role of non-canonical Tgf-β signalling in the control of epithelial progenitor proliferation and the ability of progenitors to spontaneously evade such regulatory mechanism; F.C., V.F., E.B. and D.D.F. characterized the consequences of non-canonical Tgf-β signalling evasion in organoid cultures; V.F. generated the sh-Smad4 organoid line, with the contribution of D.D.F. and M.C., carried out the pharmacological studies on Tak1/p38 MAPK signalling; V.F., A.Ala., and D.D.F. performed biochemical studies on cell cycle regulators and mediators of the type-I interferon-like response; V.F., D.D.F, S.G., G.M. and A.L. performed orthotopic transplantations; M.L. and M.G. executed Click-it mass spectrometry experiments and analysed the data; A.Be. contributed to flow cytometry; S.G. executed immunostaining experiments and contributed to image acquisition; A.Bi. contributed to the characterization of p53 function in organoids; F.B. and A.Ant. carried out and analysed chromosome painting and FISH studies; M.B., M.F. and M.L provided histopathological annotations; F.G. and A.R. performed the computational analyses; M.M.S. contributed to the interpretation of data; F.C. and V.F. assembled the figures; F.C. wrote the first draft of the manuscript; F.C., V.F. and A.L. edited the manuscript with inputs from F.B., A.Ant., A.R., M.L. and M.M.S.; A.L. acquired the main funding sources.

## Competing interests

Authors declare no competing interests.

---

**Supplementary Figure 1.**
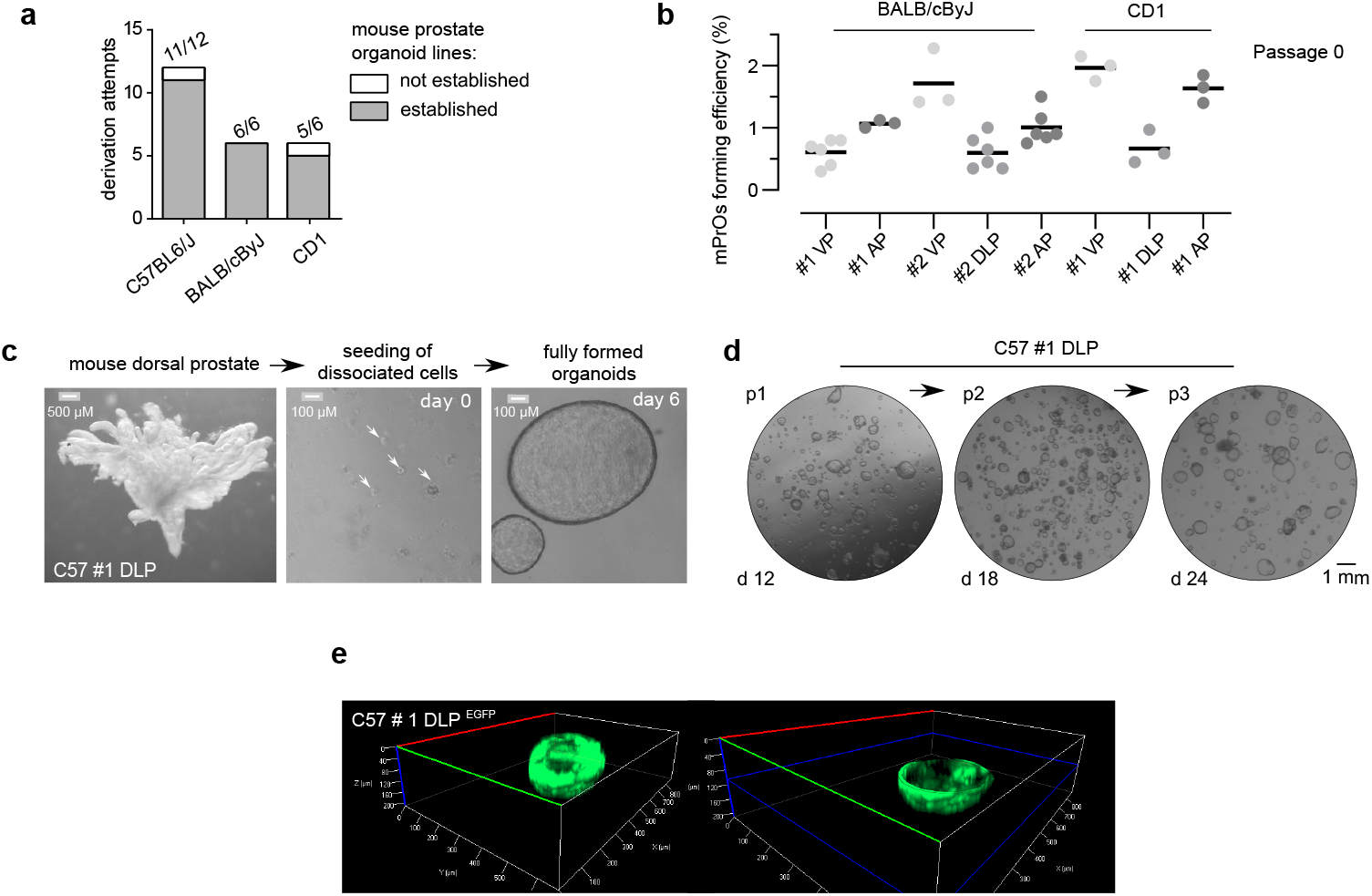
Establishment of a mouse prostate organoid biobank. **a**. Prostate organoid derivation rate from distinct mouse strains. **b**. Mouse prostate organoid forming efficiency at derivation. Data points are shown with crossing line representing mean value. **c**. Representative stereoscopic images at different stages of mouse prostate organoid derivation. **d**. Representative stereoscopic images of mouse prostate organoid cultures during subsequent passages. **e**. Representative 3D reconstruction of a fully formed mouse prostate organoid based on live epifluorescent microscopic analysis (1 μm section x 300 planes). On the right side, only the bottom half of the organoid is displayed.

**Supplementary Figure 2.**
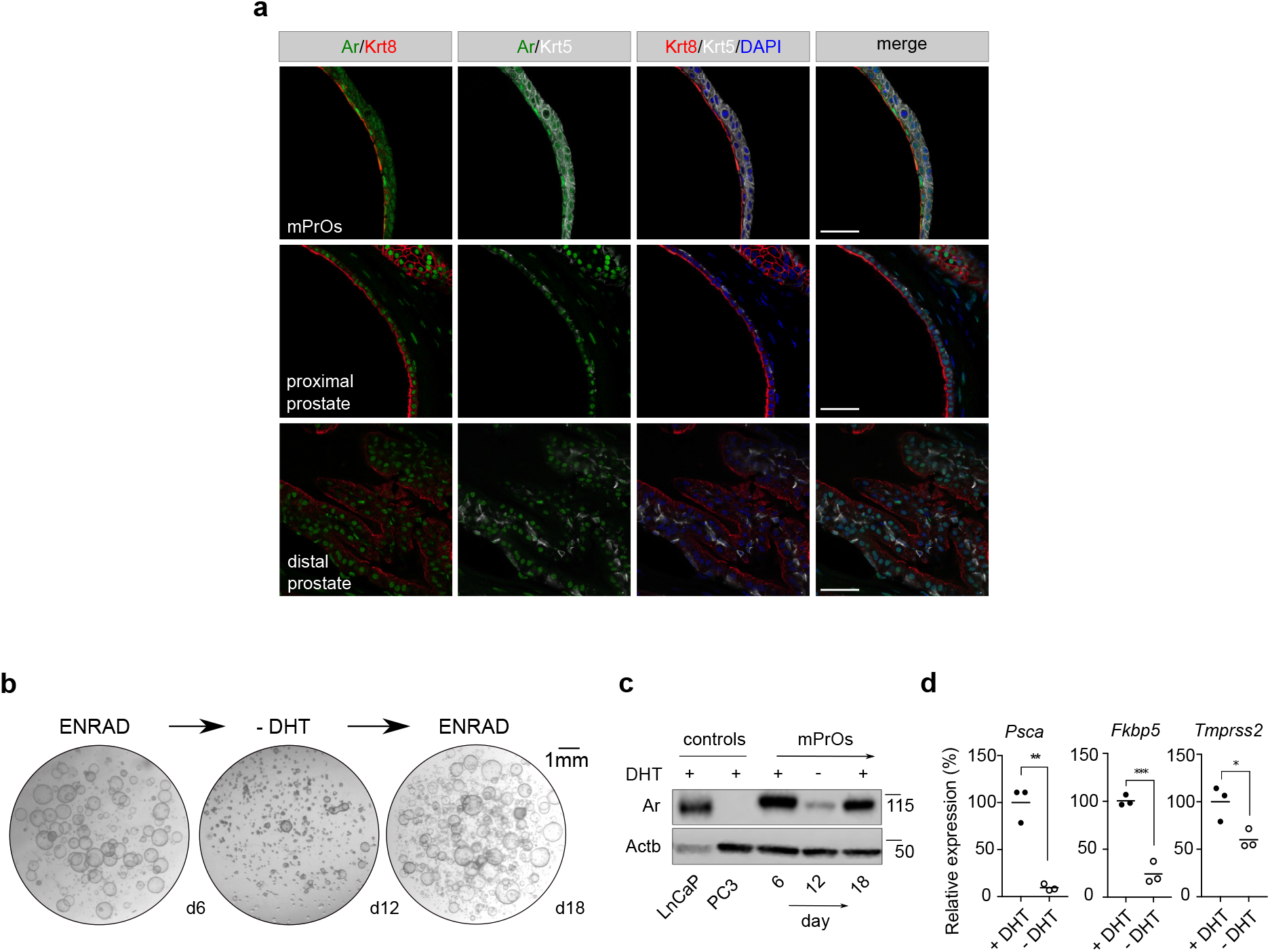
Mouse prostate organoids are dependent on androgen signalling for lumen formation - but not for survival. **a**. Immunofluorescence analysis for selected markers in mouse prostate tissue and organoid sections (scale bars = 50 μm). **b**. Stereoscopic images of mouse prostate organoids experiencing transient dihydrotestosterone (DHT) removal (androgen cycling). Lumen formation necessitates androgen signaling. **c**. Western blot analysis for Androgen receptor (Ar) expression in mouse prostate organoids experiencing androgen cycling, and control cell lines expressing (LnCaP) or not expressing (PC3) the receptor. **d**. qRT-PCR mRNA expression analysis for selected androgen-responsive genes in the presence or absence of DHT. Data points are shown with crossing line representing mean value. Student’s t-test, two-tailed, p-value * (<0.05), ** (<0.01), *** (<0.001).

**Supplementary Figure 3.**
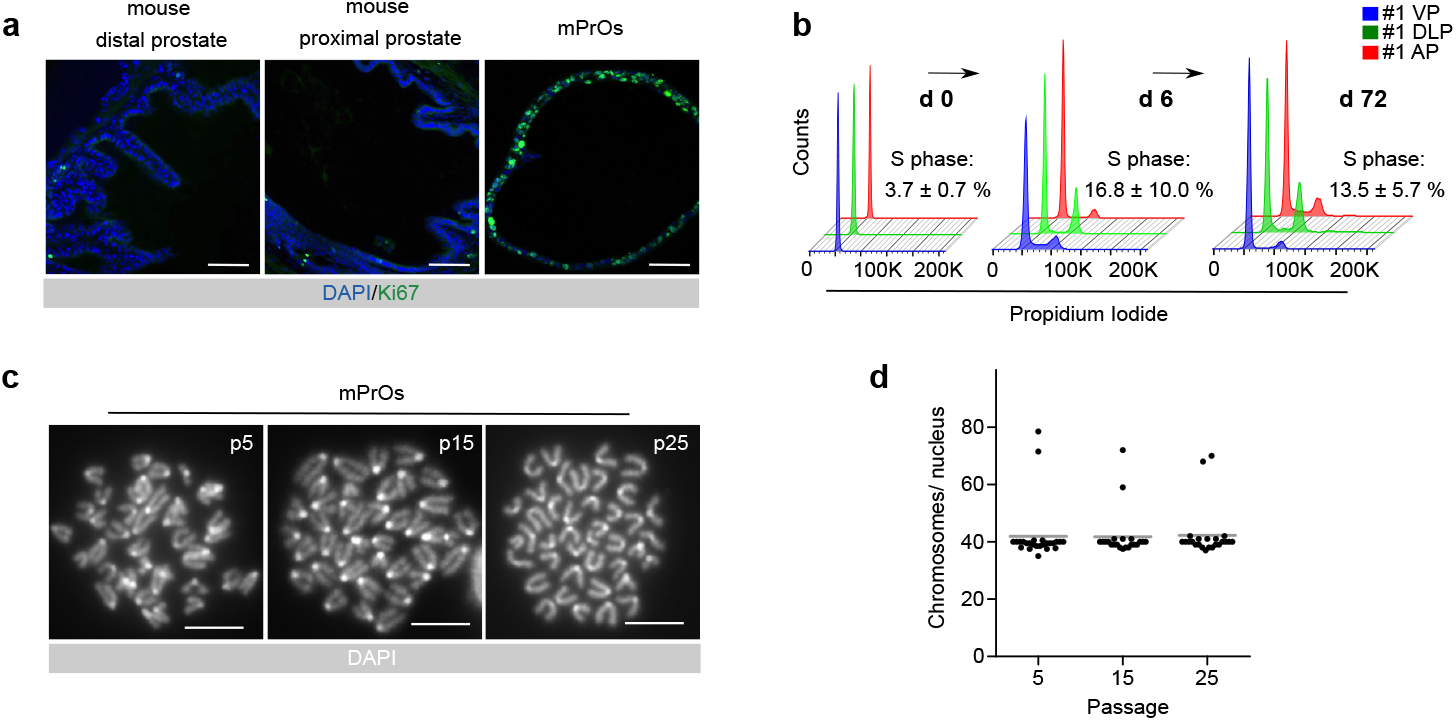
Low levels of genomic instability in fast cycling mouse prostate organoids. **a**. Immunofluorescence staining for Ki67 in mouse prostate tissue and organoid sections (scale bars = 50 μm) **b**. Flow cytometry analysis for DNA content **c**. Representative karyotypes (scale bars = 10 μm) **d**. Quantification of karyotype analysis. Data points are shown with crossing line representing mean value.

**Supplementary Figure 4.**
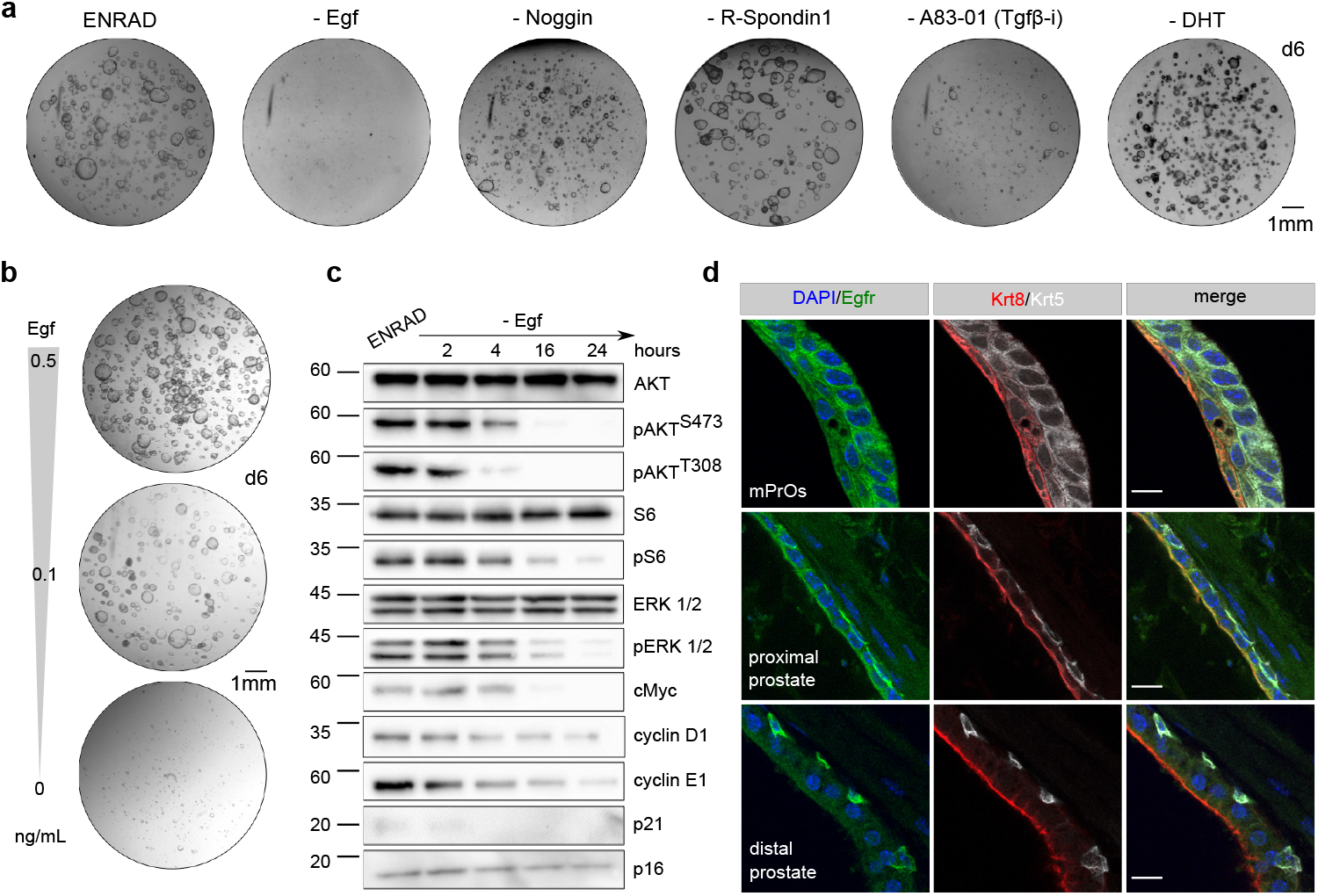
Mouse prostate organoids rely on Egf signalling for continuous proliferation. **a.** Representative stereoscopic images of mouse prostate organoid grown in complete medium (ENRAD) vs. medium depleted of individual growth factors/inhibitors. Mouse prostate organoids necessitate of both Egf and the Tgf-β inhibiitor A83-01 for continuous expansion. Within the period of observation, removal of Noggin or R-Spondin 1 has no clear consequence in culture. Please note that organoids failed to form a lumen in the absence of dihydrotestosterone (DHT). **b.** Stereoscopic images of mouse prostate organoids cultured in the presence of reduced levels (0.5, 0.1 ng/mL) or in the absence of Egf. **c.** Western blot analysis in mouse prostate organoids for selected signalling mediator and cell cycle regulator proteins. **d.** Immunofluorescence analysis for selected markers in mouse prostate tissue and organoid sections (scale bars = 10 μm).

**Supplementary Figure 5.**
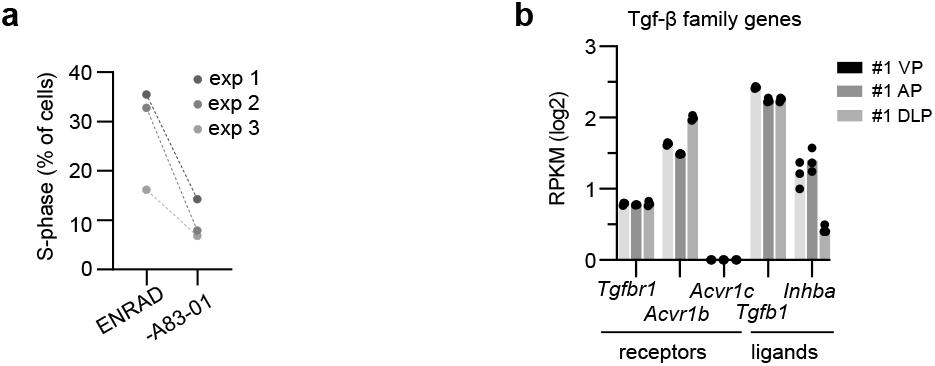
Inhibition of Tgf-β receptors is required for cell cycle progression in mouse prostate organoids. **a**. Quantification of cells undergoing S-phase based on flow cytometry (EdU *vs*. TO-PRO-3) in complete medium (ENRAD) or in the absence of A83-01 (24 hours). (Related to Fig 2a; n=3, individual data points are shown.) **b**. mRNA expression levels for selected Tgf-β family receptors and ligands. Bulk-RNAseq analysis (n=3; individual data points are shown with bar graphs representing mean value).

**Supplementary Figure 6.**
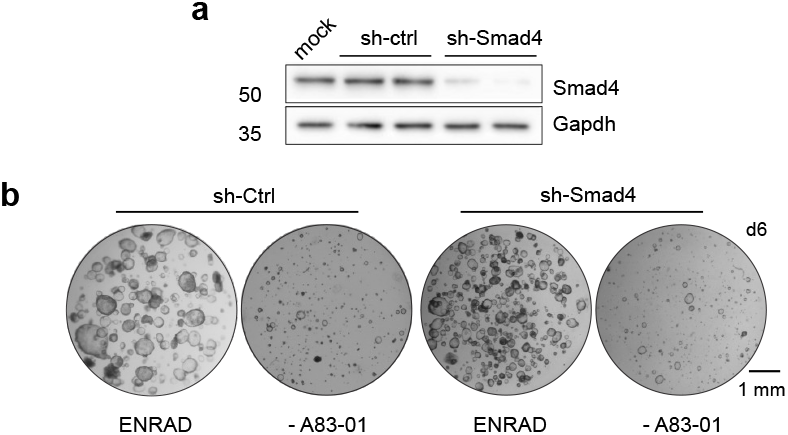
Canonical Smad signalling is dispensable for Tgf-β induced quiescience in prostate organoids. **a**. Western blot analysis for Smad4 in mouse prostate organoids upon short-hairpin RNA mediated knockdown (sh-Smad4) and in control conditions (sh-Ctrl). **b**. Representative stereoscopic images of control (sh-Ctrl) and Smad4 knocked down (sh-Smad4) mouse prostate organoids in normal growth condition (ENRAD) and following A83-01 withdrawal (-A83-01).

**Supplementary Figure 7.**
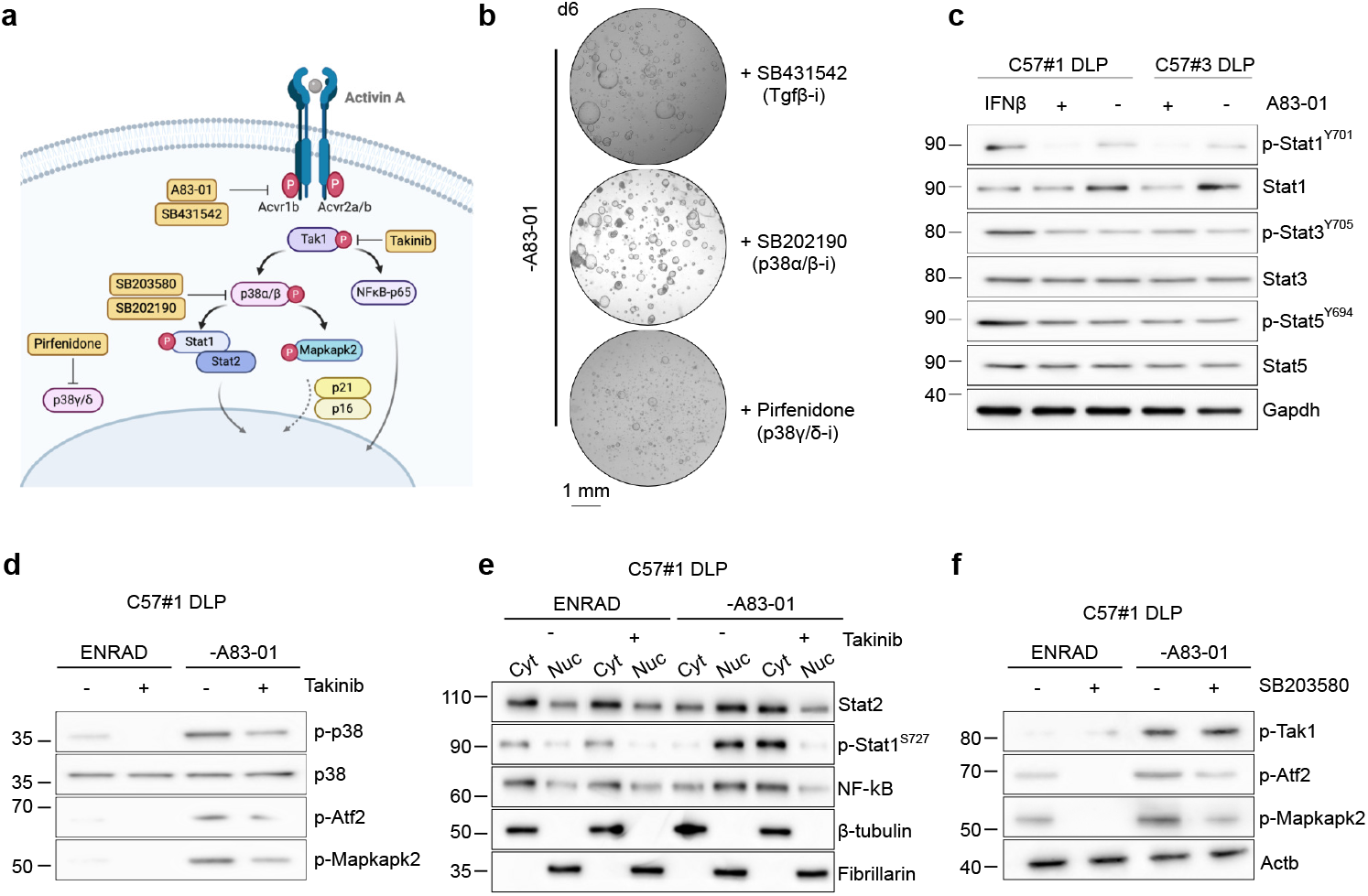
Tak1/p38 MAPK signalling activates immune- and stress-related pathways in prostate organoids. **a**. Schematic view of the non-canonical Activin A signalling pathwaty, including inhibitors used for the experiments described in this figure. **b**. Representative stereoscopic images of mouse prostate organoids grown in the absence of A83-01 and in the presence of either SB431542 (Tgfβ-receptor inhibitor; 10 μM, 6 days), or SB202190 (p38α/β inhibitor; 10 μM, 6 days), or Pirfenidone (p38γ/δ inhibitor; 10 μM, 6 days). **c**. Western blot analysis in mouse prostate organoids for canonical IFN signalling mediators upon acute A83-01 withdrawal (IFNβ; 50 ng/mL, 6 days). **d-f**. Western blot analysis in mouse prostate organoids for key components of the non-canonical Tgf-β pathway and IFN signalling upon acute A83-01 withdrawal and concomitant inhibition of either Tak1 (Takinib; 5 μM, 24 hours) (**d, e**) or p38α/β (SB203580; 10 μM, 24 hours) (**f**).

**Supplementary Figure 8.**
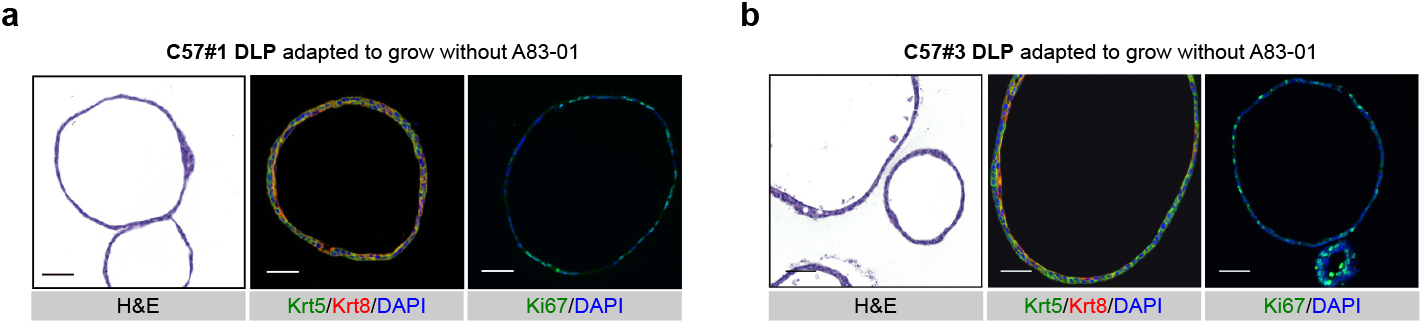
Mouse prostate organoids adapted to the absence of the Tgf-b receptor inhibitor A83-01 retain quasi-normal cellular identities and cytoarchitecture. **a-b**. Representative H&E and immunofluorescence analyses of C57#1 and C57#3 DLP mouse prostate organoids upon adaptation to A83-01 withdrawal (scale bar = 50 μm). Please note the adapted organoids retain a pattern of cytokeratins expression consistent with normal cellular identities, including basal, luminal P (lumP) and peri-uretheral (PrU)

**Supplementary Figure 9.**
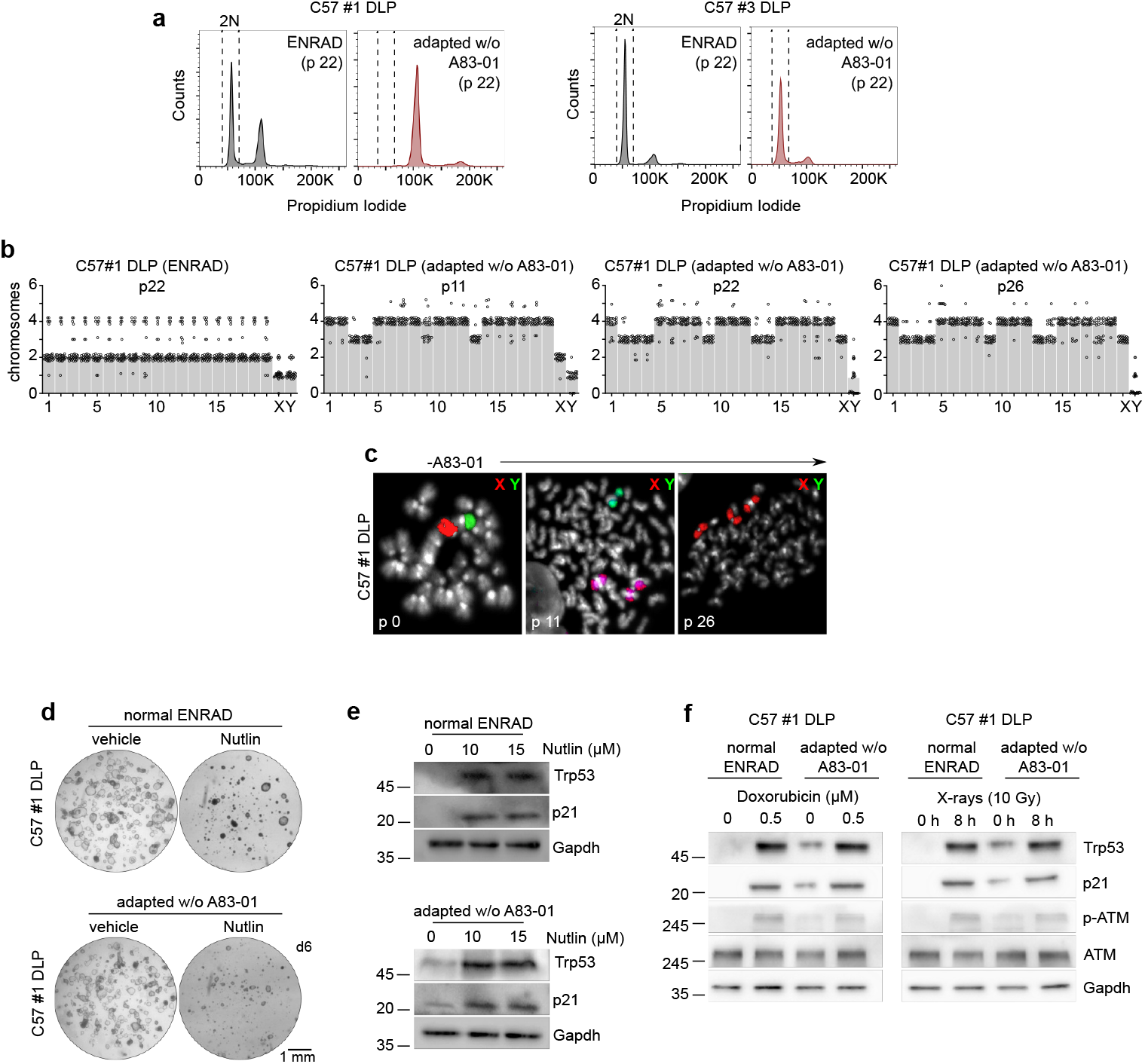
C57#1 DLP mouse prostate organoids adapted to the absence of A83-01 in culture display widespread genomic instability while retaining an intact p53 pathway. **a**. DNA content analysis in C57#1 DLP and C57#3 DLP mouse prostate organoid lines. **b**. Quantification of spectral karyotype (SKY) longitudinal analysis in C57#1 DLP mouse prostate organoids undergoing adaptation to the absence of A83-01 in culture (related to Fig 3f). During adaptation, an early genome duplication event is observed followed by selective chromosome loss and gains, and leading to subtetraploidy. **c**. Representative dual-colour FISH images for X and Y chromosomes in C57#1 DLP mouse prostate organoids during adaptation to the absence of A83-01 in culture. This analysis confirmed a progressive sex chromosome imbalance during adaptation. **d**. Representative stereoscopic images of C57 #1 DLP mouse prostate organoids treated with Nutlin, or vehicle as control. **e**. Western blot analysis of C57 #1 DLP mouse prostate organoids described in **d**. **f**. Western blot analysis in C57 #1 DLP mouse prostate organoids treated with doxorubicin or X-rays. C57#1 DLP mouse prostate organoids retain a functional p53 pathway.

**Supplementary Figure 10.**
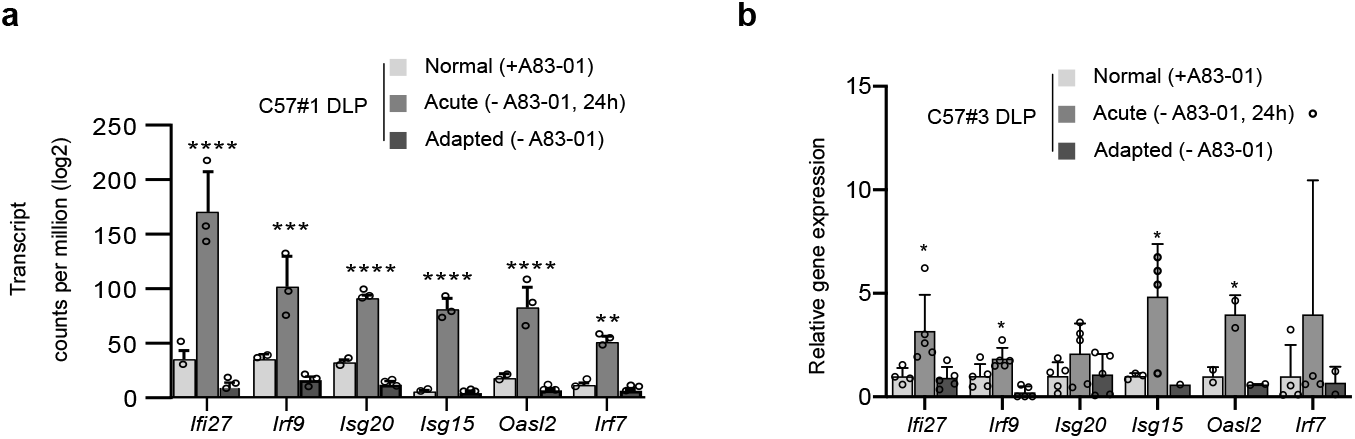
Non-canonical Activin A signalling regulates type-I interferon genes in mouse prostate organoids. **a-b**. mRNA expression levels for selected type-I IFN response genes in C57#1 and C57#3 DLP mouse prostate organoids cultured in complete medium (normal), upon acute A83-01 withdrawal (acute), and adapted to grow without A83-01 (adapted). Transcription of type-I interferon genes is enhanced upon A83-01 withdrawal and reduced upon adaptation (n=3, bulk-RNAseq analysis (a) or qPCR (b); individual data points are shown with bar graphs representing mean value and standard deviation; two-way ANOVA, Tukey’s test, p-value * (<0.05), ** (<0.01), *** (<0.001), **** (<0.0001).

**Supplementary Figure 11.**
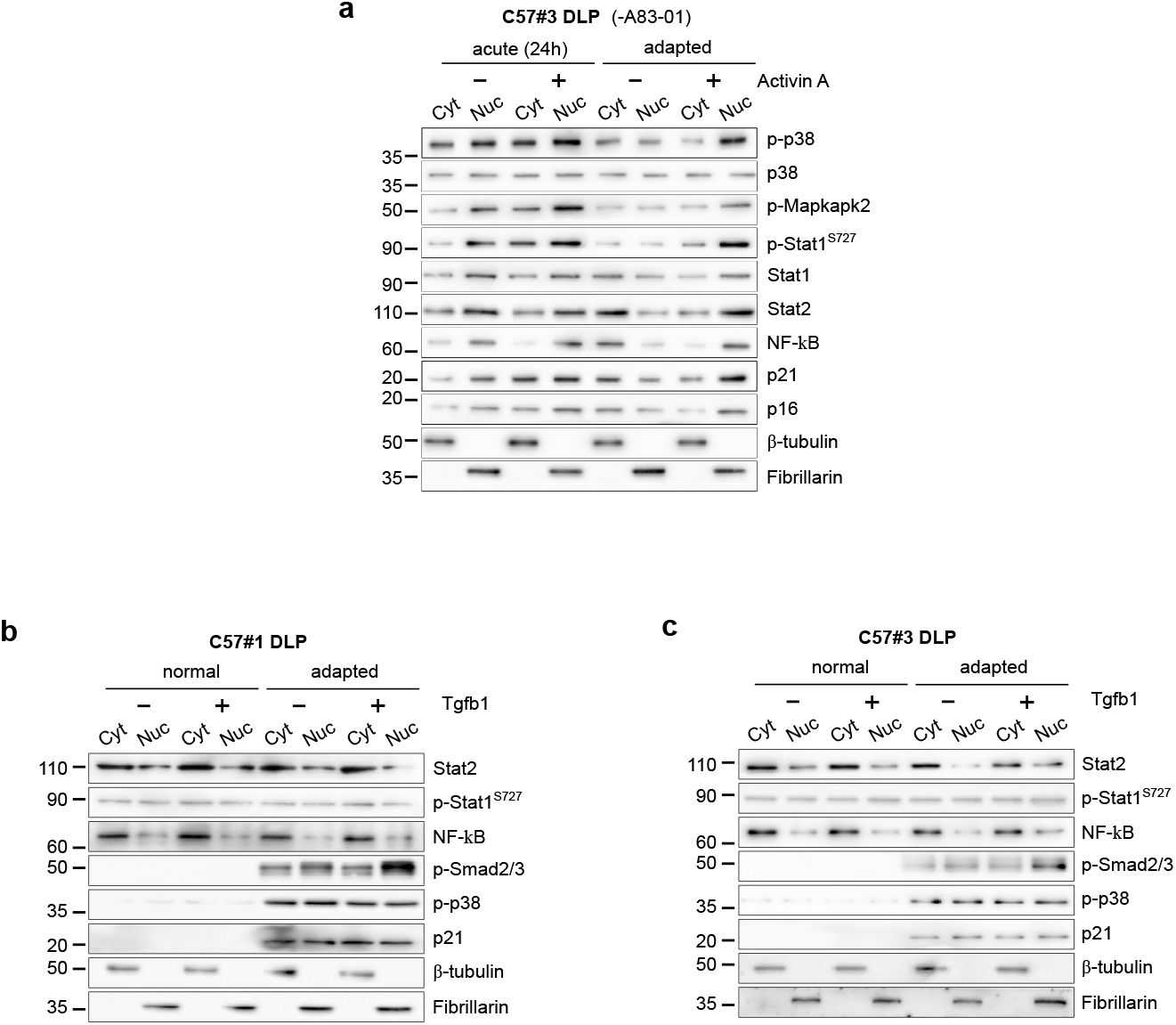
Restoration of Activin A signalling - not Tgb1 - leads to cytostasis in organoid lines (C57 #1 and #3) adapted to grow in the absence of A83-01. **a.** Western blot analysis of C57#3 DLP mouse prostate organoid line upon acute removal (24 hours) or adapted to grow without A83-01, in the presence or absence of Activin A (50 ng/mL, 24 hours). **b**. Western blot analysis of C57#1 DLP normal or adapted to grow without A83-01 mouse prostate organoid lines, in the presence or absence of Tgfb1 (500 ng/mL, 24 hours). **c**. Western blot analysis of C57#3 DLP normal or adapted to grow without A83-01 mouse prostate organoid lines, in the presence or absence of Tgfb1 (500 ng/mL, 24 hours).

**Supplementary Table 1:**
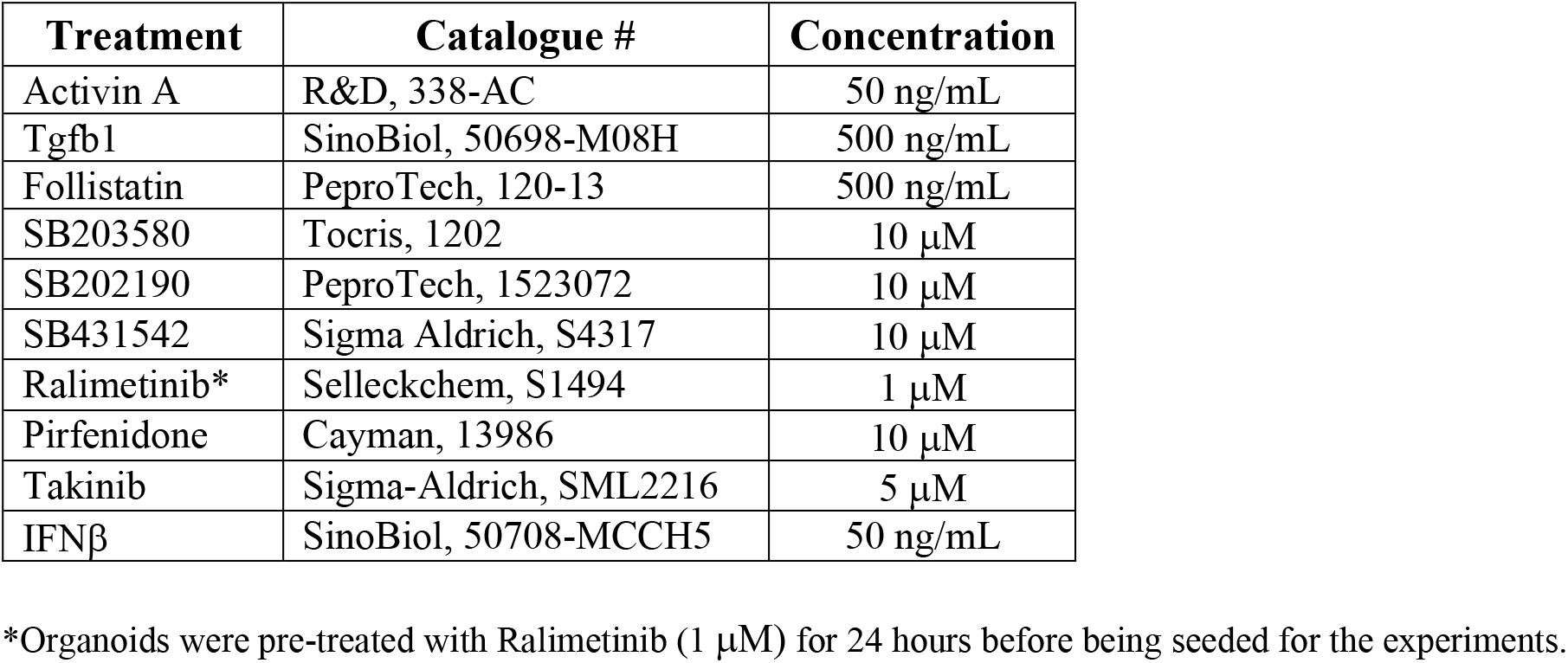
Growth factors and small molecules

**Supplementary Table 2:** Primary antibodies used for IF, IHC and flow cytometry

| **Antigen** | **Catalogue #** | **Dilution** |
| --- | --- | --- |
| Krt5 | BioLegend, 905901 | 1:500 |
| Krt8 | Merk, MAB329 | 1:200 |
| Krt7 | Abcam, ab68459 | 1:250 |
| Krt8/18 | Abcam, ab53280 | 1:1000 |
| Ki67 | eBioscience, 14-5698-82 | 1:200 |
| Egfr | Abcam, ab52894 | 1:200 |
| Acvr1b | R&D Systems, AF1477 | 1:500 |
| Ppp1r1b | Invitrogen, MA5-14968 | 1:400 |
| Ar | Santa Cruz Biotech, SC-816 | 1:500 |
| Cd24a | eBioscience, BMS17-0242-82 | 1:800 |
| Sca-1 | BioLegend, 122514 | 1:800 |

**Supplementary Table 3:**
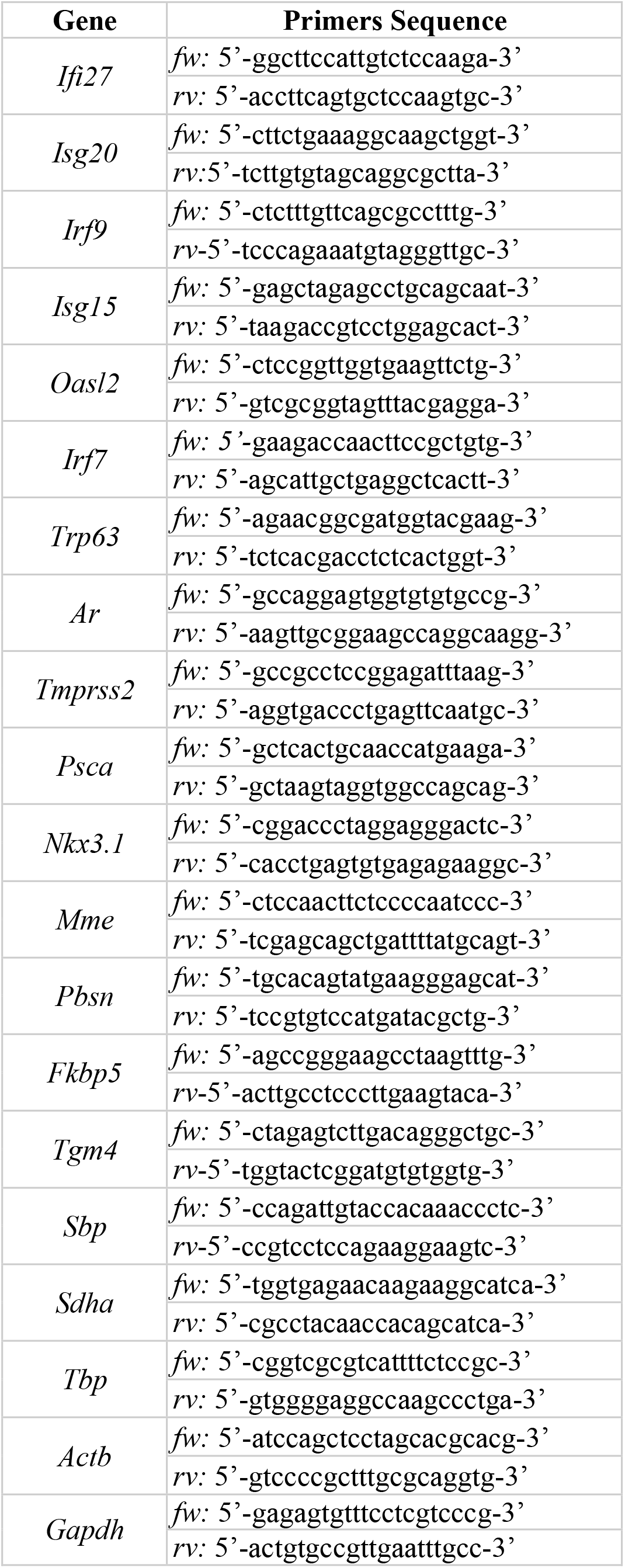
Primers used for end-point and quantitative PCR

**Supplementary Table 4:** Primary antibodies used for western blot

| **Antigen** | **Catalogue #** | **Dilution** |
| --- | --- | --- |
| β-Actin | Merk, A2228 | 1:2000 |
| β-Tubulin | Santa Cruz Biotech, sc-5274 | 1:4000 |
| Akt | CST, 9272 | 1:2000 |
| Akt phosphoSer473 | CST, 4060 | 1:1000 |
| Akt phosphoThr308 | CST, 9275 | 1:1000 |
| Ar | Santa Cruz Biotech, sc-816 | 1:500 |
| Atf2 anti-phosphoThr71 | CST, 9221 | 1:1000 |
| Atm | CST, 2873 | 1:1000 |
| Atm phosphoSer1981 | CST, 5883 | 1:1000 |
| c-Myc | Abcam, ab32072 | 1:1000 |
| Chk1 | CST, 2360 | 1:1000 |
| Chk1 phosphoSer35 | CST, 2348 | 1:1000 |
| Cyclin D1 | Abcam, ab134175 | 1:1000 |
| Cyclin E1 | CST, 20808 | 1:1000 |
| Erk | CST, 9102 | 1:1000 |
| Erk phosphoThr202/Tyr204 | CST, 4370 | 1:1000 |
| Fibrillarin | Abcam, ab4566 | 1:500 |
| Gapdh | Thermo Fisher, MA515738 | 1:4000 |
| Krt5 | BioLegend, 905901 | 1:500 |
| Krt8/18 | Abcam, ab53280 | 1:1000 |
| Mapkapk2 phosphoThr334 | CST, 3041 | 1:1000 |
| p16 | Abcam, ab211542 | 1:500 |
| p21 | Abcam, ab109520 | 1:500 |
| p38 | CST, 9212 | 1:1000 |
| p38 phosphoThr180/Tyr182 | CST, 9215 | 1:500 |
| Smad2 phosphoSer465/467 | CST, 8828 | 1:1000 |
| Smad2/3 | CST, 8685 | 1:1000 |
| Smad4 | Santa Cruz Biotech, sc-7966 | 1:1000 |
| Stat1 | BD Biosciences, 610186 | 1:1000 |
| Stat1 phosphoSer727 | CST, 9177 | 1:1000 |
| Stat1 phosphoTyr701 | CST, 9171 | 1:1000 |
| Stat2 | CST, 4597 | 1:1000 |
| Stat3 | CST, 9132 | 1:1000 |
| Stat3 phosphoTyr705 | CST, 9131 | 1:1000 |
| Stat5 | CST, 9358 | 1:1000 |
| Stat5 phosphoTyr694 | CST, 9359 | 1:1000 |
| Tak1 | CST, 5206 | 1:1000 |
| Trp53 | Abcam, ab26 | 1:500 |

## Notes

### Competing Interest Statement

The authors have declared no competing interest.

